# Force transmission is a master regulator of mechanical cell competition

**DOI:** 10.1101/2024.12.20.627898

**Authors:** Andreas Schoenit, Siavash Monfared, Lucas Anger, Carine Rosse, Varun Venkatesh, Lakshmi Balasubramaniam, Elisabetta Marangoni, Philippe Chavrier, René-Marc Mège, Amin Doostmohammadi, Benoit Ladoux

## Abstract

Cell competition is a tissue surveillance mechanism for eliminating unwanted cells and as such is indispensable in development, infection and tumorigenesis. Although different biochemical mechanisms are proposed, due to the dearth of direct force measurements, how mechanical forces determine the competition outcome remains unclear. Here, using *ex vivo* tissues and different cell lines, we have discovered an unknown form of cell competition that is regulated by differences in force transmission capabilities, favoring cell types with stronger intercellular adhesion. Direct force measurements reveal increased mechanical activity at the interface of the two competing cell types in the form of large stress fluctuations which can lead to upward forces and cell elimination. We show how a winning cell type endowed with a stronger intercellular adhesion exhibits a higher resistance to elimination while benefiting from efficient force transmission to neighboring cells. This cell elimination mechanism could have broad implications of keeping strong force transmission ability for maintaining tissue boundaries and cell invasion pathology.

## Main text

Cell competition plays a vital role in maintaining tissue health, fighting against pathogens and tumorigenesis ^1–4^. Despite these widespread and crucial implications, the fundamental principles that govern cell competition remain unclear. The elimination of loser cells can be facilitated by biochemical signals, which lead to cell death and subsequent removal ^1,2^, but various studies have also shown that cells can mechanically outcompete each other ^5,6^. The prevailing consensus is that winners compress losers, promoting loser cell’s death and removal ^3,5,6^. Different strategies such as directed migration ^7,8^, crowding ^9^, differences in cell growth ^10,11^ or homeostatic density ^12,13^ enable winning cells to apply pressure or resist to it ^7–13^. However, contradicting outcomes have emerged from studies exploring the change of cell mechanics through modulating the extracellular environment ^14–16^ or changing contractility, e.g. by overexpressing the oncogene Ras^V^^12^ in different *in vivo* and *in vitro* systems ^17–23^. Although cell competition is involved in various biological and pathological processes, a framework that integrates the role of collective mechanical interactions in cell competition is lacking. In particular, if and how cell competition is influenced by the fundamental process of intercellular force transmission is not known. Sensing, transmitting and exerting mechanical forces between cells is mediated in epithelia by the adherens junction protein E-cadherin ^24^, which is crucial for efficient intercellular mechanical coupling ^25–30^. Therefore, we conjectured that altering the intercellular force transmission through modifying the E-cadherin adhesion strength could lead to the emergence of cell competition and could strongly affect its outcome.

### Cells with higher intercellular force transmission capabilities win in cell competition

We set out to investigate if heterogeneities in intercellular adhesion strength and consequently force transmission capabilities could lead to competitive interactions. A pathological example of such molecular heterogeneities can be found in metaplastic breast cancers, a highly aggressive triple-negative breast cancer subtype presenting a therapeutic challenge ^31^. The intra-tumoral heterogeneity in force transmission capability is recapitulated by the presence of at least two sub-populations of cancer cells, epithelial and mesenchymal ^31^, with potentially varying E-cadherin expression levels in epithelial sub-population ^32^. To address how tumor cell subclones sorted and if they competed within a tumor, we cultivated patient-derived xenografts from metaplastic breast cancers and monitored their development. To focus on the role of force transmission capabilities in cell competition, we chose xenografts with a binary state in E-cadherin expression, i.e. in which E-cadherin is strongly expressed in the epithelial, but absent in the mesenchymal sub-population. The two sub-populations sorted, resulting in clusters of E-cadherin-positive epithelial cells (E-cad^+^) surrounded by E-cadherin-negative mesenchymal cells (E-cad^-^) (**Fig. 1a**). We further observed a competition between the cell types: over time, the E-cad^+^ clusters expanded at the cost of E-cad^-^ cells, removing them from the substrate (**Fig. 1b**, Video 1). This increased removal of E-cad^-^ cells was only observed when both sub-populations directly interacted (**Fig. S1 A**). We confirmed our observations using cells from a second breast cancer patient (**Fig. S1 B,C,** Video 2). These observations indeed suggests that heterogeneities in intercellular adhesion strength can lead to cancer cell competition, in which cells with increased adhesion strength win.

**Fig. 1:**
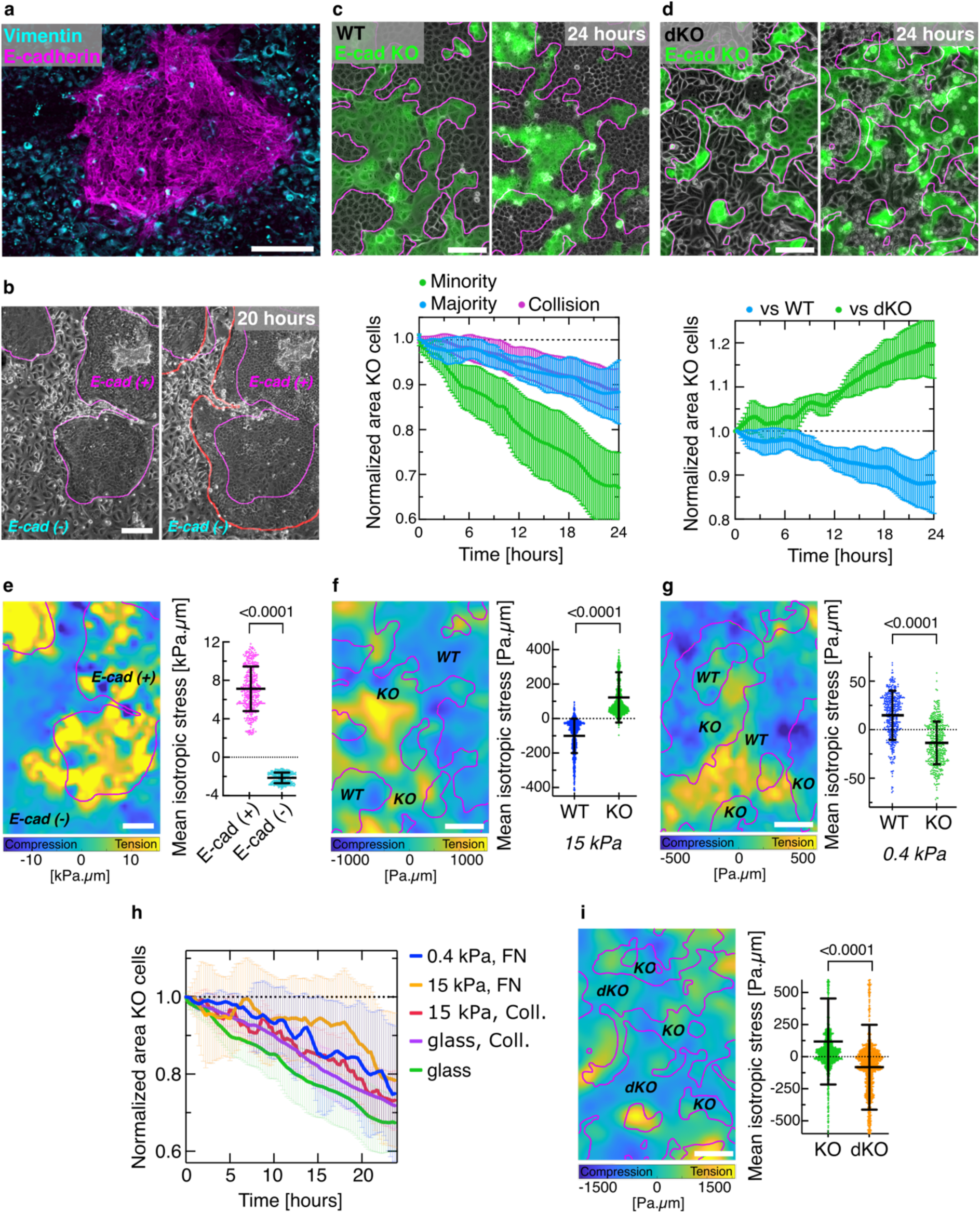
Intercellular force transmission capabilities provide a competitive advantage. **a,** Confocal image showing a monolayer of patient-derived metaplastic cancer cells. The E-cadherin positive (magenta) and vimentin positive (cyan) sub-populations sort completely. **b,** Phase contrast images of the cluster development within 20 hours. Red line shows cell clusters after 20 hours. **c,** Top: phase contrast and fluorescent images of a mixed culture of MDCK WT (grey) and MDCK E-cadherin knockout (green, fluorescently labelled with LifeAct-GFP) cells. Bottom: normalized area occupied by E-cad KO cells being in minority (green), majority (blue) or after the collision of two fully sorted populations (magenta). Collision assay realized through model wounds. Normalization to initial value, n=8 movies from N=3 independent experiments (Minority); n=6, N=3 (Majority); n=4, N=2 (Collision). **d,** Top: mixed culture E-cad KO cells (green) and E-cadherin / Cadherin-6 double knockout cells (dKO, grey). Bottom: normalized area of E-cad KO cells competing against WT (blue) or dKO (green) cells. Normalization to initial value, n=6 movies from N=3 independent experiments (vs WT); n=8, N=2 (vs dKO). **e,** Left: heatmap of the isotropic stress within metaplastic breast tumor tissue. The colormap shows compressive (blue) and tensile (yellow) stresses. The heatmap corresponds to the initial frame shown in (**b**). Right: average isotropic stress for each cell type, n=5 movies from N=2 independent experiments. **f,** Left: heatmap of the isotropic stress within the competition between WT and E-cad KO cells on 15 kPa stiffness substrates. The heatmap corresponds to the initial frame shown in (**c**). Right: average isotropic stress for each cell type, n=14 movies from N=4 independent experiments. **g,** Left: heatmap of the isotropic stress within the competition between WT and E-cad KO cells on soft substrates (370 Pa). Right: average isotropic stress for each cell type, n=7 movies from N=2 independent experiments. **h,** Normalized area occupied by E-cad KO cells on substrates with different stiffnesses and surface coatings (uncoated glass in green, glass coated with collagen in magenta, 15 kPa PDMS coated with fibronectin (FN) in orange, 15 kPa PDMS coated with collagen in red and 370 Pa PAA coated with FN in blue). E-cad KO cells are in minority. They are under tension on stiff and under compression on soft substrates. Normalization to initial value, n=8 movies from N=3 independent experiments (glass); n=7, N=1 (glass, Coll.); n=10, N=3 (15 kPa, Coll.); n=10, N=3 (15 kPa, FN) and n=10, N=3 (370 Pa). **i,** Left: heatmap of the isotropic stress within the competition between E-cad KO and dKO cells on 15 kPa stiffness substrates. The heatmap corresponds to the initial frame shown in (**d**). Right: average isotropic stress for each cell type, n=13 movies from N=2 independent experiments. All datapoints represent the mean value of all isotropic stresses within one field-of-view of one frame. P-values from unpaired t-test. All magenta lines show initial cell clusters. All error bars show the standard deviation. Scale bars 200 µm (**a, b, e**); 100 µm (**c, d, f, g, i**).

To investigate the role of intercellular adhesion in cell competition more systematically, we turned to competition between two other cell types: we lowered the adhesion strength of MDCK epithelial cells by knocking out E-cadherin (E-cad KO). In pure cultures, E-cad KO cells showed no signs of reduced cell viability and due to the presence of cadherin 6, they still form mechanically active junctions, although of lower strength ^33^. Mixing E-cad KO and WT cells, we observed that the populations sorted ^33,34^ and that the E-cad KO cells were outcompeted by the WT cells (**Fig. 1c**). Because both cell types showed a similar cell density (**Fig. S2 A**), we used population area change to estimate cell losses. To quantify population areas, E-cad KO cells were expressing lifeAct-GFP. These cells lost against normal WT cells as well as WT cells expressing lifeAct-mCherry (**Fig. S2, B**), excluding an impact of lifeAct expression on the competition. Importantly, E-cad KO cell loss was independent of cell ratios, as they also lost when in majority (**Fig. S2 C**). To better control the boundary between the two cell types, we developed a collision assay of two migrating cell populations ^18,35^ which led to the same competition outcome (**Fig. S2 C**, Video 3). To assess if and how E-cad KO cells compete against cells with even further reduced cell-cell adhesion, we mixed them with MDCK E-cadherin/cadherin 6 double knockout cells (dKO) which cannot form any adherens junctions ^36^. The previously losing E-cad KO cells won and outcompeted the dKO cells (**Fig. 1d**, Video 4). To modulate force transmission strength through the expression levels of E-cadherin, we then mixed WT and E-cadherin overexpressing ^37^ (E-cad OE) cells. WT cells were eliminated by these cells with even further increased cell-cell adhesion (**Fig. S3 A-D**). To generalize our findings, we performed similar experiments with another epithelial cell line which originates from breast tissue, MCF10A cells, mixing WT and E-cad KO cells ^38^. The E-cad KO cells were also eliminated by the WT cells, both in mixed cultures (**Fig. S3 E**) and in collision assays (**Fig. S3 F**). Taken together, experiments across diverse cell types show, without exception, that cells with relatively stronger adherens junctions always win in cell competition, including patient-derived tumors and various epithelial cell lines.

### Winning cells can be under tensile or compressive stresses

Cell elimination can be governed in epithelia by compressive stresses ^39–41^. As force transmission within tissues is mainly regulated through adherens junctions ^25,27,42^, we first reasoned that stronger intercellular adhesion could allow winning cells to collectively exert compressive stresses on the losing cells, in line with current consensus described in the literature ^7–9,11–13,17^. Unlike previous studies, our experimental setup provides additional information that includes direct access to intercellular stresses using Bayesian Inversion Stress Microscopy ^41,43,44^. In the patient-derived tumor cultures, we observed that the winning E-cad^+^ cells were under high levels of tension and the losing E-cad^-^ cells under compression (**Fig. 1e**), in agreement with their strong differences in stiffness (**Fig. S4 A,B**) and exerted traction forces (**Fig. S4 C**). However, to our surprise in mixtures of MDCK WT and E-cad KO cells, winning WT cells were under compression and losing E-cad KO cells were under tension (**Fig. 1f**). This non-intuitive, unanticipated, observation is contrary to established models ^7–9,11–13,17^. We confirmed this result with the collision assay in which we controlled temporally the establishment of the contact between the two cell types. The mechanical state of WT cells switched from tensile during migration to compressive after the collision with the E-cad KO cells (**Fig. S5 A**). Similar results were obtained using another force interference method independent of traction forces and based on cell shape obtained from labelling tight junctions ^45^ (**Fig. S5 B,C**). They were further confirmed by laser ablation experiments (**Fig. S5 D**), were WT cells showed a negative (Video 5, compression) and E-cad KO cells a positive recoil (Video 6, tension) (**Fig. S5 E**). Although E-cad KO cells were on average under tension, local regions remained under compression (**Fig. 1f**). Thus, we wondered whether E-cad KO cells were eliminated preferentially at these local compressive regions. Assessing the isotropic stresses locally (**Fig. S6 A,B**) prior to cell elimination revealed that E-cad KO cells were under tension before and during the elimination process (**Fig. S6 C**). This confirms that the competition outcome is independent of local compressive regions. To further compare this mechanism to previously established cell competition scenarios that include loser cell death^7,9,10^, we investigate the fate of eliminated cells by labelling dying cells with annexin V. We observed that 70% of E-cad KO cells were eliminated alive and only later died due to their extraction from the tissue and thus the absence of adhesion ^46^ (**Fig. S6 D,E**). Furthermore, we inhibited apoptosis using a pan-caspase inhibitor, which did not change the competition outcome (**Fig. S6 F**). Together, this data shows that the cell elimination mechanism is independent of loser cell death. Moreover, since cell competition based on biochemical signaling usually leads to cell death ^1,2^, live cell extrusion strongly supports a cell elimination mechanism based on mechanical forces.

To investigate other competition scenarios, we changed the mechanical environment of all cells using softer substrates (370 Pa) to lower the cell-substrate adhesion ^44^ (**Fig. S7 A**) and exerted tractions (**Fig. S7 B**). Under such conditions, E-cad KO cells were now under compression and the WT cells under tension (**Fig. 1g**) but the competition outcome remained the same, i.e. WT cells won independently of substrate composition or stiffness (**Fig. 1h**). We further measured stresses in the competition between E-cad KO and dKO cells and observed the same pattern of tension-compression with winners, E-cad KO cells, under tension and losers, dKO cells, under compression (**Fig. 1i**). Overall, we show that compression-induced cell loss can indeed explain the outcome of different competition scenarios. However, the direct measurement of intercellular stresses challenges this established consensus that winners always squeeze out losers. Demonstrating that cells can be under compression and still win suggests that other, still unknown mechanisms must be governing the cell competition outcome.

### Elimination of E-cad KO cells cannot be explained by any established mechanism

To understand why the E-cad KO cells were losing despite being under tension, we first ruled out previously conjectured mechanisms. For instance, differential cell growth could impact cell competition ^11,12,10^, but both cell types exhibited identical fractions of mitotic cells (**Fig. S8 A**) and similar growth rates (**Fig. S8 B**) in pure and mixed cultures. Cells with higher homeostatic density can have a competitive advantage ^13,7^, but cell competition emerged at cell densities well below the homeostatic density of both cell types (**Fig. S8 C**). Quantifying the rates of cell elimination, both cell types showed similar extrusion rates in pure cultures, and the rates increased with time and cell density (**Fig. S8 D**). In mixed cultures, however, the rate of extrusion was strongly increased for E-cad KO cells compared to pure cultures, and independent of cell density, while the rate of extrusion for WT cells remained comparable to pure cultures (**Fig. 2a, Fig. S8 D**). This demonstrates that the predominant elimination of E-cad KO cells in mixed cultures was not due to cell-intrinsic processes but resulted from their collective interactions with WT cells. Previous reports on the role of cell mechanics in cell competition have conjectured that a relative increase in cell-substrate adhesion ^8,16^ and cell stiffness ^12,17^ provides a competitive advantage. Furthermore, E-cadherin based adherens junctions have shown to be mechanosensitive, affecting various aspects of cell and tissue mechanics ^26,33,47–49^. Thus, we assessed how the decrease of cell-cell adhesion strength (**Fig. S9 A,B**) had globally affected E-cad KO cell mechanics. The cells capacity to form tight- or desmosome junctions was not changed (**Fig. S5 B; Fig. S9 C**). This underlines that the mechanical link between the cells is only weakened. In mixed cultures, E-cad KO cells exerted significant larger traction forces on the substratum than their WT counterparts (**Fig. S9 C**) and showed a striking increase in focal adhesion size (**Fig. 2b, Fig. S9 D**). Using surface indentation, we measured a significant increase in E-cad KO cell stiffness compared to WT cells for pure and mixed cultures (**Fig. S9 E**), most likely due to their more prominent actin-based contractile phenotype ^33^ (**Fig. 2c**). These observations demonstrate that a cell population’s ability to generate increased forces and exert them on competing cells does not necessarily provide a competitive advantage: Loser cells can exhibit stronger cell-substrate adhesions and higher stiffness, which explains the state of tension in eliminated E-cad KO cells, but make their elimination even more puzzling, contradicting the proposed cell-substrate and cell stiffness advantage ^8,12,16,17^. Finally, contact-dependent cell-cell signaling could lead to cell elimination independent of mechanical forces^1,2^. However, we observed that E-cad KO cells were eliminated not only at the interface of the two populations, but also more than one cell row away from it (**Fig. S10 A**, Video 7). In conclusion, having tested multiple possibilities, we ruled out the applicability of previously reported mechanisms in explaining the outcome of WT and E-cad KO cell competition, suggesting that a new, hitherto unknown, mechanism must be at play.

**Fig. 2:**
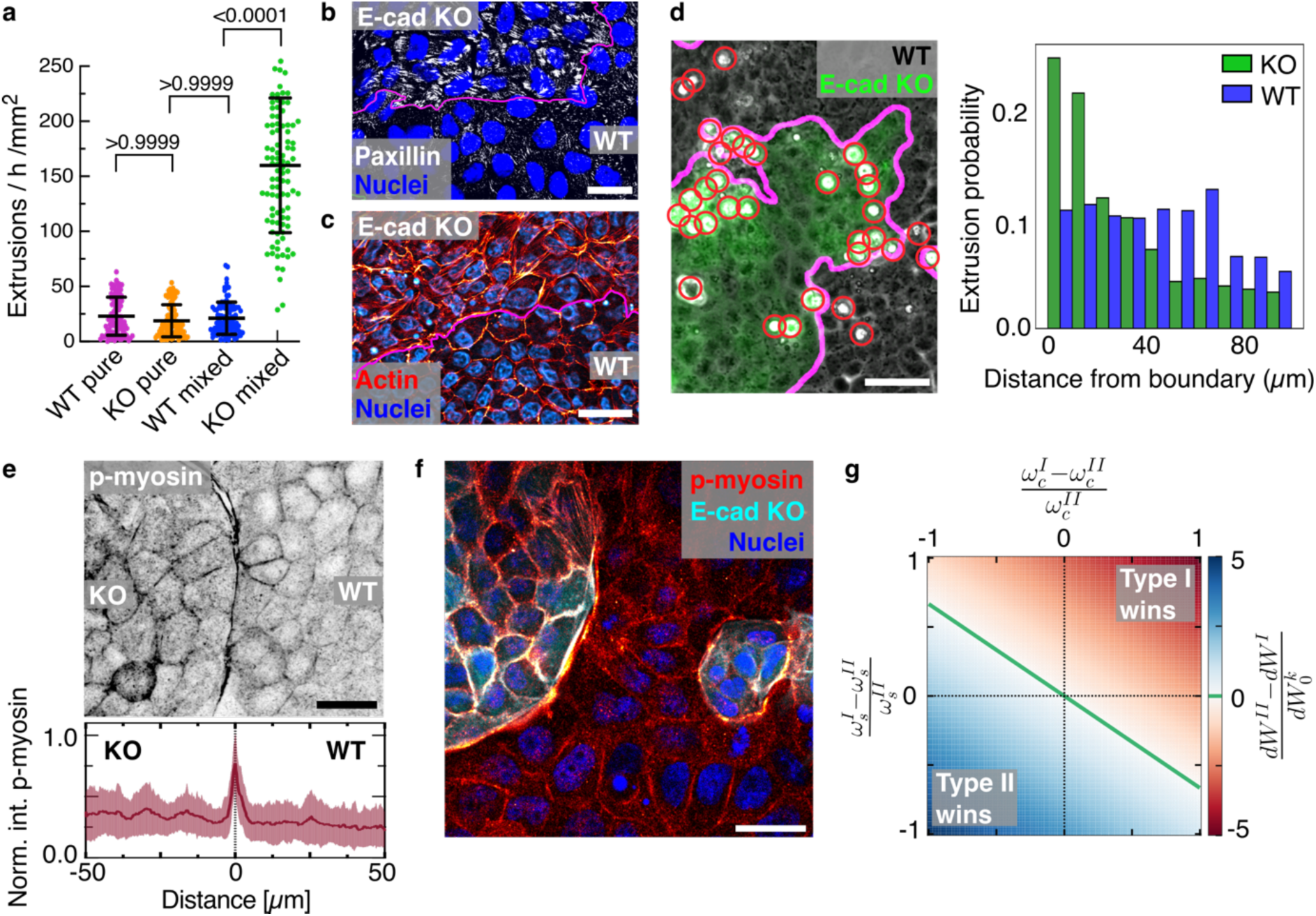
E-cad KO cells are eliminated at a mechanically active interface. **a,** Extrusion rates calculated by counting all extrusions within 1-hour intervals normalized to the area occupied by each cell type during the time interval. E-cad KO cells are in minority in the mixed culture. P-values from Kruskal-Wallis test corrected for multiple comparisons (Dunn’s test). Extrusions quantified for n=100 intervals representing 8 movies from N=2 (pure cultures) or N=3 (mixed cultures) independent experiments. **b,** Relative mechanical properties of E-cad KO cells. Example images of Focal Adhesions (Paxillin, white) and **c,** the actin cytoskeleton (red, maximum projection). **d,** Left: representative phase contrast and fluorescent image of MDCK WT and E-cad KO cells (green). Red circles indicate extrusions. Right: probability distribution of cell extrusion regarding the distance from the interface for WT and E-cad KO cells. Cells have a typical diameter of 10-15 µm. n=14729 KO extrusion, n=11031 WT extrusion, from N=4 independent experiments. **e,** Top: inverted grayscale image of phospho-myosin at the tissue interface. Bottom: normalized intensity of phospho-myosin obtained from line plots crossing the tissue interface. Lines were centered at the interface (dashed line, distance=0), indicated by fluorescent signal from E-cad KO lifeAct-GFP (not shown), averaged from n=24 measurements from N=2 independent experiments. **f,** Example fluorescent image of pluricellular actomyosin cables forming in WT cells observed in coculture. Cables are enriched in phospho-myosin (red, intensity-coded) and form along islands of E-cad KO cells (cyan). **g,** Phase diagram showing the work required to eliminate cells. X-axis shows the difference in cell-cell adhesion. Y-axis shows the difference in cell-substrate adhesion. Color code indicates difference in work, i.e. indicates winning and losing. The top left region shows that cells with relative high cell-cell adhesion can win despite lower relative cell-substrate adhesion. All error bars show the standard deviation. Scale bars 50 µm (**d**); 25 µm (**c**, **e**, **f**); 10 µm (**b**).

### Cell elimination is localized at a mechanically active interface

To further explore the preferred elimination of E-cad KO cells, we investigated the spatial distribution of extrusion events. Previous studies on mechanical cell competition proposed that loser cells get eliminated in the bulk of the cell cluster, where compressive stress is the highest ^6^. Moreover, increased contractility at tissue interfaces can impact cell elimination during development ^20,22^, but its role remains elusive ^21^. We found that the losing E-cad KO cells were preferentially eliminated near the interface, while the WT cell extrusions showed a relatively homogenous distribution (**Fig. 2d**). The WT cells did not show an increased cell density at the interface (**Fig. S10B**); thus, E-cad KO extrusions were independent of local WT densities. Importantly, neither the free edge of isolated E-cad KO monolayers (**Fig. S10 C**) nor an interface of a confined E-cad KO layer with a rigid passive fence (**Fig. S10 D**) recapitulated the predominant localization of extrusions at the interface. This suggests that the preferred elimination of E-cad KO cells is triggered by the active interface that emerges between the two tissues with contrasting mechanical properties. Accordingly, the shared interface of E-cad KO and WT cells was strongly enriched in phosphorylated actomyosin in both cell types (**Fig. 2e,f, Fig. S11 A**), indicating an increased mechanical activity there. LifeAct and phospho-myosin colocalized (**Fig. S11 A**). Using live cell imaging, we observed a polarization in actin accumulation only at the shared interface the cell types first collided (**Fig. S11 B**), which underlines the increased interface force generation. In this vein, we extended our analysis to the patient-derived tumor cultures. As in MDCK cells, the losing E-cad^-^ cells were extruded at the tissue interface (**Fig. S11 C**), where increased actomyosin activity was observed (**Fig. S11 D**). MDCK WT cells could even form pluricellular actomyosin cables at areas of high negative curvature (**Fig. 2f**). We hypothesized that pluricellular formation of actomyosin cables might help WT cells in efficiently removing small E-cad KO clusters through purse-string mechanisms as observed in wound closure ^24^. However, E-cad KO cells got eliminated at both regions of positive and negative curvatures (**Fig. S12**). Thus, such cables cannot be a dominant factor here. Independently of curvature, the enrichment in active myosin could generate a mechanical barrier, which prevents mixing of cell types through which they might confine each other ^20^. Together, the correlation between cell elimination and high mechanical activity suggests a critical role of this active interface in determining the outcome of cell competition. We then postulated that cell types endowed with different mechanical properties might react differently to this increased interface activity. To predict how energetically costly it is to eliminate each cell type, we considered a simplified analytical model for energetic requirements of cell elimination: at the interface, two competing cells pull and push on each other leading to deformations of the cells. Thus, the work done on each cell type to deform and eventually eliminate it can be expressed in terms of the energies associated with cell-substrate and cell-cell adhesions strengths, and cell stiffness (see Methods). The energy required to remove a cell can be simply estimated as the work required to deform the cell from a cylindrical shape to a cone-like shape and then rounding it up to a sphere upon cell removal (**Fig. S13**). Comparing the work required to eliminate the competing cells as a function of the difference in their cell-cell adhesion strength demonstrates that the cell type with a higher cell-cell adhesion could require more work to be eliminated, even if the other type has a higher cell-substrate adhesion (**Fig. 2g**). This simple energetic argument shows that the energy barrier for elimination is higher for cells with strong cell-cell adhesion. As such, this minimal model, does not consider where the energy required for elimination comes from and therefore does not tell anything about the mechanism driving the elimination. To bridge this gap, we next employ a more detailed, cell-based model that resolves individual cells, their interactions, and mechanics.

### A computational model reveals stress fluctuations lead to interfacial cell elimination

To understand how the active interface affects mechanical competition and why strong cell-cell adhesion presents a competition advantage therein, we turned to physical modeling of 3D cell monolayers ^50^. Our model is based on a multi-phase fields approach that accounts for both passive and active interactions of deformable cells in three-dimensions (3D). These interactions include cell-cell and cell-substrate adhesion strengths that are considered explicitly and tuned independently (see **Fig. S14** for model schematics). This enables modulating force transmission capability and its effect on the competition outcome while providing access to the out-of-plane 3D stress components that govern the removal of cells from a monolayer (see Methods). A cell extrusion is captured in the model without any explicit threshold, or external artificial means to favor one. Once the out-of-plane forces acting on a cell overpower the forces keeping it in the monolayer and on the substrate, a cell extrusion occurs. In this vein, the collective behavior of cells, e.g. cell extrusion and height fluctuations, emerge from solving the dynamics associated with translation and interface relaxation of each cell (see Methods) (**Fig. S15**). To best represent the experimental conditions, we modeled collision assays of two model cell types (**Fig. 3a**, Video 8): model wild type (mWT), and model E-cad KO (mE-cad KO) defined based on cell-cell adhesion differences (lower for mE-cad KO) and/or cell-substrate adhesion contrast (higher for mE-cad KO). In agreement with the experimental observations, mE-cad KO cells, with a higher cell-substrate adhesion and a lower cell-cell adhesion relative to mWT cells, were eliminated at the interface (**Fig. 3b**). To understand why E-cad KO cells are eliminated at the interface, we quantified the fluctuations in stress fields *via* susceptibility ^51,52^, which is defined as as *χ* = *N* × [⟨σ^2^⟩ − ⟨σ⟩^2^], where ⟨⟩ indicates expectation and *N* is the number of data points corresponding to σ, σ^2^ fields. The susceptibility of isotropic stress field primarily due to in-plane fluctuations (**Fig. S16**), 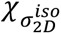, and linked to out-of-plane component of stress tensor, *σ_zz_*, peaked at the interface of mE-cad KO and mWT cells, a consequence of contrasting physical properties of the cell types considered (**Fig. 3c**). At the same time, in-plane stress fields exhibited a weaker correlation in mE-cad KO cells relative to mWT cells, suggesting a muted ability to transmit stresses (**Fig. 3d**). Additionally, the out-of-plane component of the stress field, *σ_zz_*, near the interface exhibited a pronounced localization in mE-cad KO cells relative to their mWT counterparts (**Fig. 3e**), particularly in the tensile region (**Fig. 3f**). To further investigate the link between in-plane fluctuations and out-of-plane stress localization, we considered a series of simulations where cell-substrate adhesion contrast is kept constant, while the contrast in cell-cell adhesion is increased, by reducing the cell-cell adhesion strength of mE-cad KO cells. Interestingly, in-plane susceptibility near the interface decreased (**Fig. 3g**) while the location of extrusion events shifted away from the interface (**Fig. 3h**) as the contrast in cell-cell adhesion is reduced. These results suggested that higher in-plane fluctuations led to more extrusions of mE-cad KO cells near the interface. To understand why, we focused on stress transmission away from the interface.

**Fig. 3:**
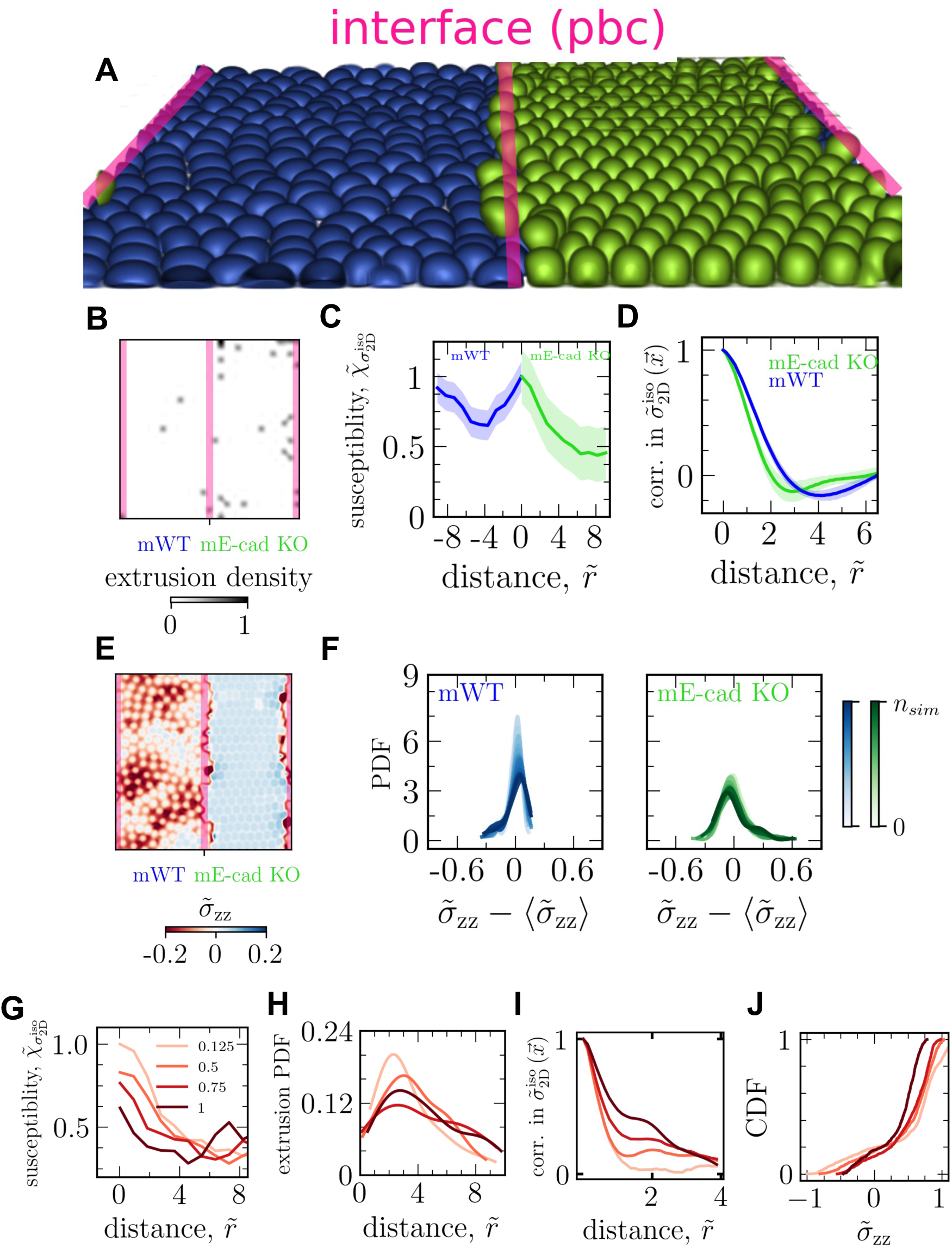
Computational model reveals the role of high fluctuations at the active interface in determining the outcome of cell competition. **a,** Example simulation snapshot with mE-cad KO cells (Green) losing to mWT cells (Blue) at the interface (red lines), keeping in mind the periodic boundary conditions (PBC). **b,** Extrusion density map representing spatial distribution of extrusion events, corresponding to the simulation in (**a**). **c,** Susceptibility of two-dimensional, i.e. in-plane, isotropic stress field and the out-of-plane component of stress tensor normalized by the maximum value in mE-cad KO cells for each, as a function of distance from the interface. The distance is normalized by the initial cell radius. The data corresponds to the simulation in (**a**). **d,** Spatial correlation of the in-plane, i.e. two-dimensional, isotropic stress for each cell type corresponding to the simulation in (**a**). **e,** Out-of-plane stress component field, normalized by maximum value of in-plane compression. **f,** Probability density function (PDF) for fluctuations in out-of-plane stress component, normalized by maximum value of in-plane compression for each cell type near the interface within the distance of four times cell radius on each side. The color shades capture the temporal evolution of the PDFs where is the total number of time steps. **g,** Susceptibility of in-plane isotropic stress field for mE-cad KO cells for fixed cell-substrate adhesion and various cell-cell adhesion strengths (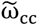) normalized by the value for the lowest cell-cell adhesion at the interface. **h,** Extrusion PDFs corresponding to (**g**). **i,** Spatial correlations corresponding to coarse-grained in-plane isotropic stress fields averaged, ensemble and temporal, centered around an extruding cell in a square domain of eight times cell radius for fixed cell-substrate adhesion and varying cell-cell adhesions corresponding to (**g**). **j,** cumulative distribution functions (CDF) corresponding to the average out-of-plane stress fields normalized by maximum in-plane compression around an extruding cell, showing higher localization for lower cell-cell adhesion: the peak shifts to the left and becomes less tensile as cell-cell adhesion increases.

We noted a more persistent susceptibility away from the interface i.e. a relatively smaller difference in susceptibility near the interface and further from it, by increasing cell-cell adhesion (**Fig. 3g**). More importantly, characterization of the spatial correlation of averaged, in-plane, isotropic stress fields prior to and at the onset of extrusion (**Fig. 3i**) showed that these fields became more correlated as the cell-cell adhesion of mE-cad KO cells are increased, signaling a more efficient transmission of mechanical information. Indeed, inspection of the out-of-plane component of the averaged fields, *σ_zz_*, around extrusions show higher localization due to ineffective stress transmission by mE-cad KO cells with low cell-cell adhesion (**Fig. 3j**), resulting mE-cad KO cells to extrude near the interface. In summary, the *in silico* study showed (i) the emergence of an actively fluctuating interface due to differences in cell-cell adhesion strengths and (ii) weakening cell-cell adhesion hindered the flow of mechanical information away from this active interface, manifesting in less correlated stress fields. This explained why mE-cad KO cells are eliminated at the interface. Unable to transmit the high in-plane isotropic stress fluctuations away from the interface, mE-cad KO cells seek relief by localizing stresses out-of-plane and potentially extruding as mWT cells expand into their domain.

### Experiments confirm strong interface stress fluctuations driving cell elimination

To verify these predictions, we first assessed experimentally the susceptibility of mechanical stresses and found the same striking increase of stress fluctuations at the interface, which correlates with the localization of E-cad KO extrusions (**Fig. 4a**). As expected, the increase of fluctuations at the interface was also found in the substrate displacement and in the traction forces (**Fig. S17 A**). Additionally, in line with simulation predictions of enhanced fluctuations at higher cell-cell adhesion difference, we observed even stronger interface fluctuations in the primary tumor sample where the difference in cell-cell adhesion is higher relative to MDCK cells (**Fig. S17 B**). Besides differences in cell-cell adhesion, we hypothesized that high cellular activity is required for high stress and traction fluctuations. To investigate cellular behavior at the interface, we assessed the dynamics of the actin cytoskeleton. E-cad KO cells were highly active and extended several µm long protrusions below surrounding WT cells (**Fig. 4b**, Video 9). This dynamic protrusion activity demonstrated an increased cellular motility of E-cad KO cells at the interface, in line with increased traction fluctuations. We inhibited protrusion formation using CK666. E-cad KO cells did not lose any more (**Fig. S17 C**), which additionally supports that interface fluctuations are crucial for their elimination. Furthermore, increased mechanical activity could lead to increased stress fluctuations. To reduce the mechanical activity, we treated MDCK cells with blebbistatin. It globally inhibits actomyosin-generated cellular forces, which might have variable effects on the entire cell population. Blebbistatin disrupted the interface between the cells, evident by a reduced interface convexity (**Fig. 4c, Fig. S18 A, B**). Importantly, blebbistatin decreased stress magnitudes (**Fig. S18 C**), leading to cell relaxation ^53^. Due to their increased contractility, this relaxation is relatively stronger in E-cad KO cells, increasing the area of single cells (**Fig. S18 D**), and leading to an E-cad KO domain increase after blebbistatin addition (**Fig. S18 E**). The reduction of cellular forces led to a striking drop in stress fluctuations (**Fig. 4d**), which correlated with a significant reduction in the global extrusion rate of E-cad KO cells, while the extrusion rate for WT cells remained the same (**Fig. S18 F**). These experiments confirm the emergence of increased stress fluctuations at mechanically active tissue interfaces and indicate that maintaining the active interface is required for WT cells winning.

**Fig. 4:**
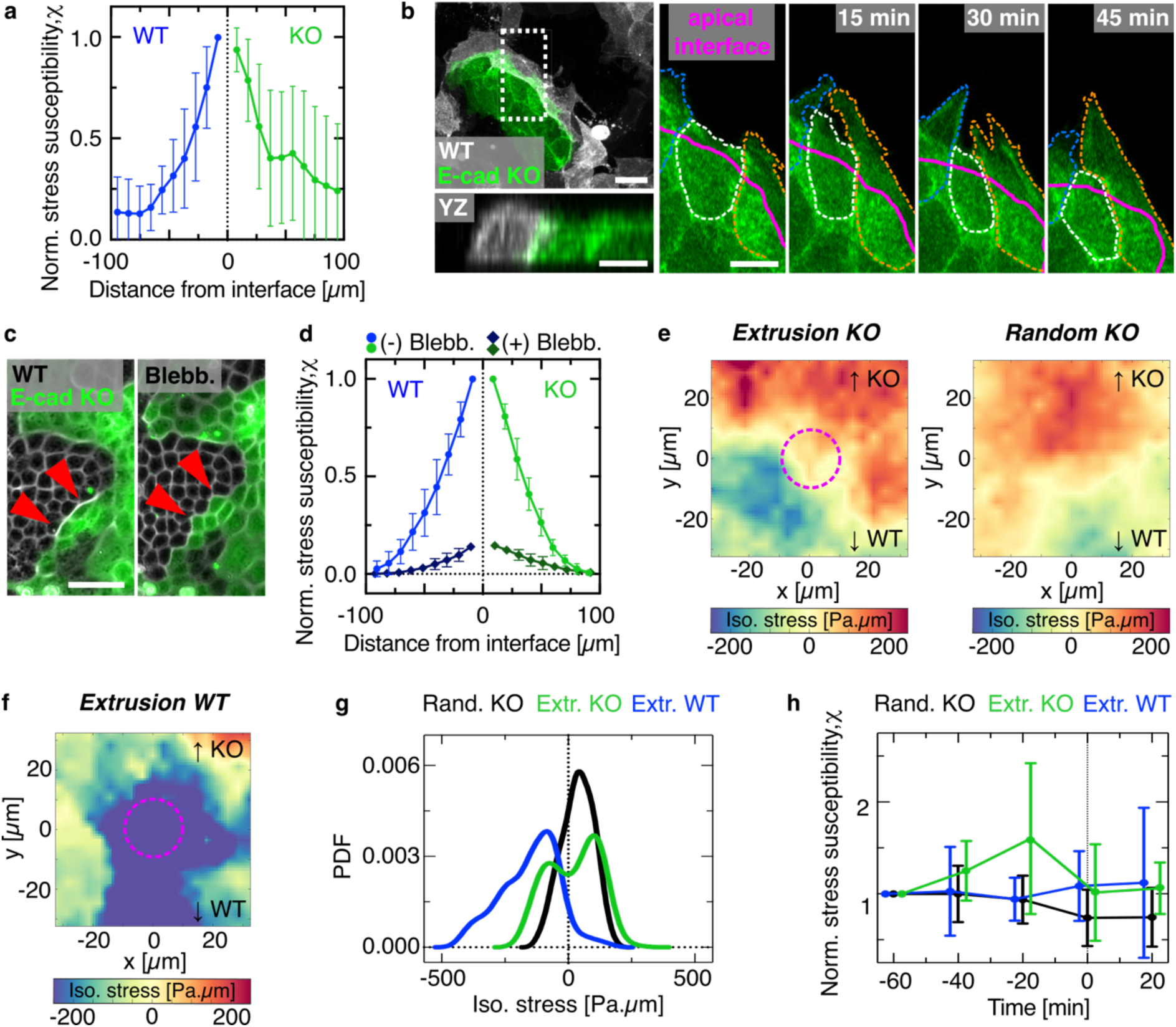
E-cad KO cells are eliminated through increased stress fluctuations. **a,** Isotropic stress susceptibility as a function of the distance from the interface for each cell type normalized to the maximum value. n=18 movies from N=5 independent experiments including mixed cultures and collisions. **b,** Snapshots of actin dynamics at the interface. Maximal projections and side view of MDCK E-cad KO LifeAct-GFP (green) and MDCK WT LifeAct-Ruby (white). Right: Zoom-in on maximal projection of E-cad KO cells protruding below WT cells. Apical interface indicated by magenta line drawn based on WT LifeAct-Ruby signal (not shown). **c,** Phase contrast and fluorescent image of the boundary (red arrows) before (left) and 2 hours after 20 µm blebbistatin addition (right). **d,** Isotropic stress susceptibility versus distance from the interface for each cell type, before (dots) and after (rectangles) addition of blebbistatin and normalized to the highest value. n=10 movies from N=2 independent experiments. **e,** Left: ensemble average heatmap of the local isotropic stress before E-cad KO extrusions. Extrusions considered were within a 30 µm distance from the interface and stresses were averaged up to 40 min before automated detection of the extrusion, excluding the timepoint of completed extrusion. Stress fields were oriented based on E-cad KO fluorescent signal so that the KO side is on top and the WT side on the bottom (see Methods). Right: average heatmap of the isotropic stress within a square of size 30 µm around random position in E-cad KO occupied area within a 30 µm distance from the interface. n=798 extrusions (E-cad KO), n=750 KO random positions from 4 independent experiments. **f,** Average heatmap of the isotropic stress before WT bulk extrusions. n=741 extrusions from N=4 independent experiments. **g,** Probability density functions of the average isotropic stress distribution before extrusion detection (WT cells bulk elimination, blue, KO cells interface elimination, green, and random KO interface position, black) corresponding to (**e, f**). **h,** Temporal evolution of the mean isotropic stress susceptibility before (t<0) and briefly after an extrusion event. t=0 indicates timepoint of automated detection of extrusion. Random position of E-cad KO cells at the interface: black, E-cad KO elimination: green, WT cell elimination: blue. Susceptibility is averaged within a square of size 60 µm around one extrusion event. Normalized to initial value, n=726 extrusions (E-cad KO), n=1050 KO random positions and n=334 extrusions (WT) in n=6 movies from N=2 independent experiments. All error bars show the standard deviation. Scale bars 50 µm (**c**); 10 µm (**b**).

To further explore the relationship between interface stress fluctuations and cell elimination, we assessed the local stress fields before cell extrusions close to the interface by computing the ensemble average stresses up to 40 min before the extrusion event. The stress field around cell extrusion events in E-cad KO cells exhibited high values of both compressive and tensile stresses (**Fig. 4e, left**) whereas the one at random positions at the interface were under lower values of tensile stresses (**Fig. 4e, right**). This indicates that E-cad KO cells experienced increased fluctuations of stresses prior to their elimination at the interface. By contrast, the stress field around extruding WT cells, which showed no preference for being eliminated at the interface (see **Fig. 2**), was exclusively compressive (**Fig. 4f**). These findings are confirmed by the distribution of isotropic stresses, which showed a much wider range and more extreme values of both compressive and tensile stresses for E-cad KO cells destined to extrude compared to E-cad KO cells at random positions (**Fig. 4g**). We next analyzed the temporal evolution of local stress fluctuations up to 60 min prior to extrusion and compared them to fluctuations at random positions along the interface. While the fluctuations at random positions and for WT cells remained relatively stable, we observed a strong and significant increase starting 40 min before E-cad KO cell extrusion events (**Fig. 4h, Fig. S19 A**). Post removal, these fluctuations returned to the initial level (**Fig. 4h**). Together, these different mechanical signatures of cell elimination point towards different cell elimination mechanisms: WT cells are extruded through high compressive stresses ^39–41^. In contrast to that, we found another cell elimination mechanism as E-cad KO cells are eliminated at the interface through increased stress fluctuations. In the competition between two cell types, this latter mechanism based on stress fluctuations can be dominant and governs the outcome.

### Collective stress transmission and deformation prevent cell elimination

Cells at the interface were subjected to increased stress fluctuations, but only the ones with lower intercellular adhesion were eliminated. Therefore, we reasoned that high intercellular adhesion must endow the winning ones with mechanisms to resist stress fluctuation-mediated elimination. The computational model predicted more efficient stress transmission to neighboring cells, preventing the localization of out-of-plane stresses in winning cells (**Fig. 3d-f**). Indeed, WT cells showed a significantly increased stress correlation length compared to E-cad KO cells (**Fig. 5a, b**). This confirms a more efficient transmission of mechanical stress to neighboring cells for WT cells. The observation of multicellular actomyosin cables between WT cell (see **Fig 2**) but not between E-cad KO cells (**Fig. S19 B**) supports these measurements. Furthermore, we reasoned that the proposed mechanism of stress fluctuations at the interface should be reflected in deformations and changes in cell shape. To this end, we assessed the cell height and the cell-cell adhesion area of WT and E-cad KO cells. Although both cell types normally have the same height (**Fig. 5c, top**), cell shapes fluctuated near the interface and particularly, WT cells could morph into columnar shape (**Fig. 5c, bottom**). The differences in cell shapes on the collective level were most striking in the collision assay, where the WT cell deformation started from the boundary and extended over multiple cells into the bulk. However, the E-cad KO cells did not deform collectively and cells away from the interface remained flat (**Fig. 5d, e**). The WT cells strongly deformed within the first 12 hours following collision (**Fig S19 B,C**), which correlated well with the increasing E-cad KO extrusion rate (**Fig S19 D**). Moreover, when surrounded by E-cad KO cells, islands of WT cells could collectively sustain high deformation and drastic cell area fluctuations over hours without being extruded (**Fig. 5f**). Such cell shape changes thus increased the intercellular contact zone between WT cells, allowing them to further increase their adhesive energy to better resist stress fluctuations. Notably, some doublets of WT cells were eliminated by E-cad KO cells, suggesting that isolated WT cells cannot propagate stresses and lose their advantage (**Fig S19 E**). The mirror situation revealed that islands of E-cad KO cells did not undergo such strong deformations and released stresses through cell elimination by extrusion (**Fig. 5g**). Together, these experiments confirmed enhanced stress transmission in winning cell types and further show that keeping strong intercellular adhesion allows the winning type to resist elimination through substantial cell shape deformations.

**Fig. 5:**
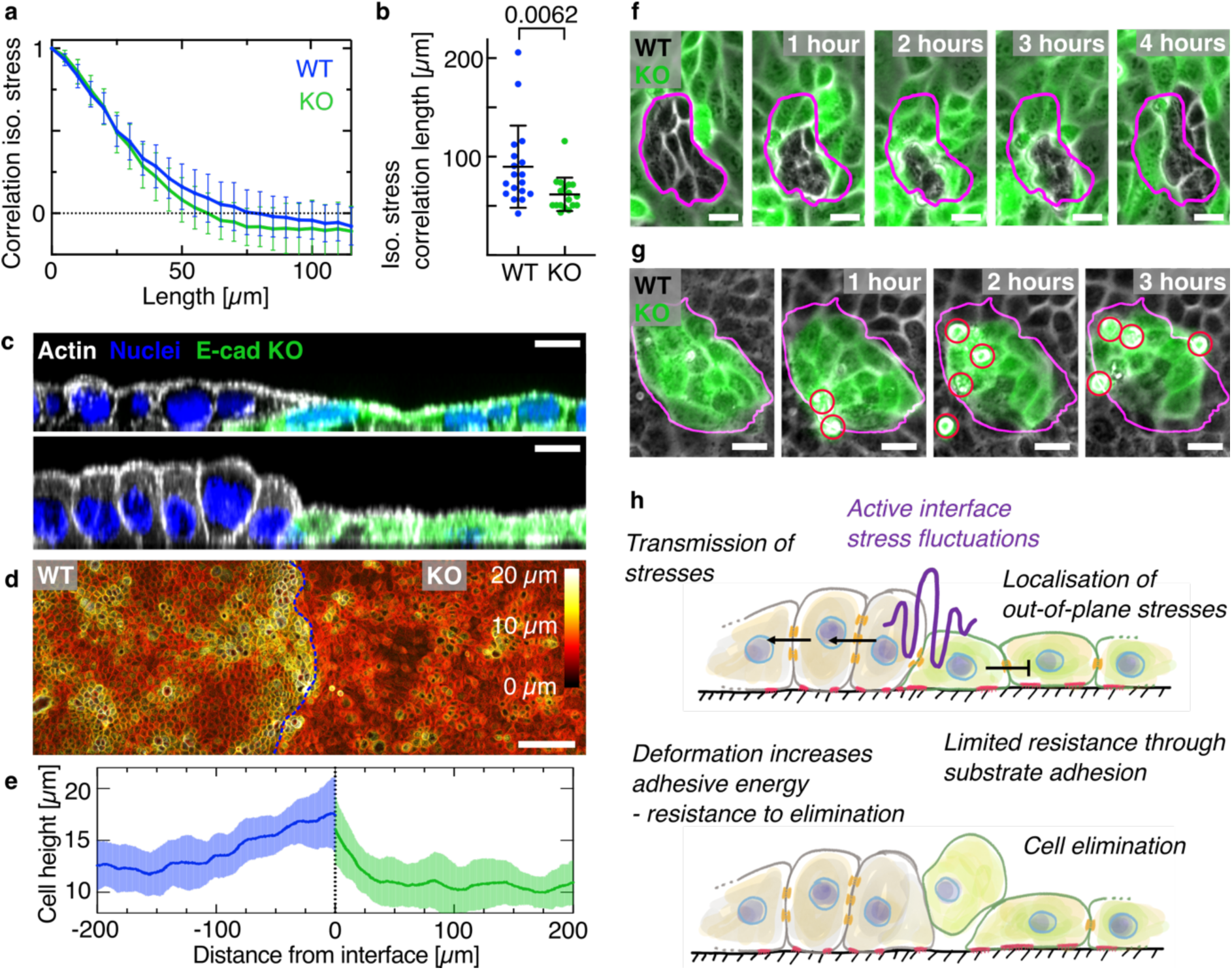
Increased intercellular adhesion endows cells with increased resistance to elimination. **a,** Spatial auto-correlation of the isotropic stress for each cell type. The zero-crossing indicates the correlation length. n=18 movies from N=5 independent experiments. **b,** Average correlation length of the isotropic stress for each cell type. P-value from unpaired t-test. **c,** Deformation of WT cells close to the interface. Confocal image showing side views of actin (white), lifeAct-GFP in E-cad KO cells (green) and the nuclei. Top: Example cell height in mixed culture. Bottom: Infrequent increased height for WT cells. **d,** Color-coded height projection of actin signal in collision assay. **e,** Quantification of cell height in collision. E-cad KO cells exhibits a faster decrease of cell height with distance from the interface compared to WT. n=15 positions from N=2 independent experiments. **f,** Brightfield and fluorescent (E-cad KO, green) images of WT cells getting compacted without cell elimination. **g,** E-cad KO cells responding to island compaction through cell elimination. **h,** Sketch of proposed mechanism. All error bars show the standard deviation. Scale bars 100 µm (**d**); 20 µm (**g**); 10 µm (**c**, **f**).

## Discussion

Here, we discover that differences in force transmission capability directly determine the outcome of mechanical cell competition, in which cells with stronger intercellular adhesion are exclusively winning. Because no previously described cell elimination mechanism could explain our observations, we propose a new one based on combining simulation and experiments. We note that a possible contribution of secreted, extracellular factors to cell elimination ^1^ cannot be excluded completely. Comparing stress patterns across multiple competition scenarios demonstrates that cells with increased force generation are under tension, which compresses the other cell population. Because these stress patterns can not predict winning and losing, we propose that the force transmission, rather than the force generation capability governs the competition outcome. Thus, our proposed mechanism is independent of loser cell compression ^7–9,17,20^ and differences in growth rate or homeostatic density ^11–13^. Increased fluctuations of isotropic stresses emerge at active interfaces between tissues with different mechanical properties. These high fluctuations in local stress fields near the interface, if not transmitted efficiently by the front-line cells to the rest of the collective, localize and induce out-of-plane stresses, akin to Poisson effect in elasticity, which can lead to cell elimination. In scenarios where cells with heterogeneities in force transmission capability compete against each other, intercellular adhesion provides a generic winning strategy since it enables winning cells to withstand higher fluctuations of stresses than losing cells. Thus, unlike other forms of mechanical cell competition such as directed migration towards losing cells ^7,8,15^, our findings unveil an alternative mechanism based on active resistance to elimination through a reinforcement of intercellular adhesion. Indeed, cells with higher intercellular adhesion can transmit stresses more efficiently to neighboring cells which prevents the localization of elimination-promoting out-of-plane stresses. In addition, increased intercellular adhesion allows collective cell shape changes into a columnar shape, which increases the mechanical threshold required for elimination through further increasing the adhesive energy. By contrast, cells with relatively lower intercellular adhesion are eliminated through the localization of high stress fluctuations at the interface and an overall limited resistance to out-of-plane stresses (**Fig. 5h**). Our conclusions are based on a physical model, only relying on the effect of mechanical imprints. Thus, if similar mechanical imprints are given, this proposed framework could have important implications for different biological processes beyond cell competition. Since it does not rely on loser cell death, it could play a role in organizing tissues during morphogenesis. Force transmission differences could be involved in maintaining tissue boundaries and, thus, functionality in homeostasis. In the skin, for example, loser cells are expelled apically through an unknown mechanism, and failed competition leads to deteriorated barrier function ^55^. The reduction of intercellular adhesion has been associated with metastasis for a long time ^56^. Adding to these mechanisms, increased stress fluctuations at the interface of tumoral and normal tissues could also play a role in invasion mechanism or promote metastasis, if tumoral cells are eliminated alive ^6,57^. Moreover, as suggested by our experiments using patient-derived tumor xenografts, this mechanism of cell competition could be acting within tumors with heterogeneities in their cell-cell adhesion strength. While this study is focused mainly on binary expression of adhesion molecules, further studies need to address heterogeneity of Cadherin expression levels, which are present in other breast cancers ^32^. Mechanical cell competition might change the fate of cells, *i.e.* promote invasion and subsequent metastasis of sub-populations. Thus, it will be exciting to further explore the role of this form of cell competition in tissue sculpting and different pathologies.

## Acknowledgments

We thank the members of the “Cell Adhesion and Mechanics” team for helpful discussion. We acknowledge the ImagoSeine core facility of Institute Jacques Monod, member of France-BioImaging (ANR-10-INBS-04) and IBiSA, with support of Labex “Who Am I”, Inserm Plan Cancer, Region Ile-de-France and Fondation Bettencourt-Schueller. We thank Tien Dang for establishing different cell lines. We thank Susana Godinho for sharing MCF10A E-cad KO cells. We thank Wang Xi for providing soft hydrogels. We thank Matteo Spatuzzi for help with data annotation. We thank Philippe Marcq for help with BISM.

## Author contributions

A.S., S.M., L.A., R.M.M., A.D. and B.L. designed the research. A.S. and L.A. performed most and analyzed all the experimental data. S.M. performed phase field simulations. C.R. performed tumor xenograft experiments. V.V. performed numerical simulations. L.B. provided movies of WT/E-cad KO competition. E.M. provided patient tumor samples. P.C. provided materials. A.S., S.M., L.A., R.M.M., A.D. and B.L. wrote the manuscript. All authors read the manuscript and provided input on it.

## Data and materials availability

Data, materials and image analysis code are available upon request.

## Supplementary Materials

- Materials and Methods
- Figures S1 to S19
- Movies S1 to S9

