## Supplementary material for "Force transmission is a master regulator of mechanical cell competition": Methods, materials and Supplementary figures

**The PDF file includes:**

Materials and Methods
References
Supplementary Figures Fig. S 1 to 19

**Other Supplementary Materials for this manuscript include the following:**

Movie S1 to S7

### Materials and Methods

Substrate preparation

*Polyacrylamide gel preparation*

370 Pa soft polyacrylamide gels were prepared as described previously (*1*). In brief, glass coverslips were cleaned in an ethanol bath, sonicated for 1 minute and dried at 80 °C for 15 minutes afterwards. The coverslips were treated with high power plasma in a plasma cleaner for 10 min. Then, they were soaked in a silane solution consisting of 2% 3-(trimethoxysilyl)propylmethacrylate (catalog no 440159, Sigma Aldrich) and 1% acetic acid in ethanol for 30 min. The silanized coverslips were rinsed with ethanol, dried at 80 °C for 1 hour and stored at room temperature.

Plasma cleaned glass coverslips or plasma cleaned glass-bottom imaging dishes (Fluorodish, WPI) were incubated with 5% fibronectin (Sigma) in PBS and dried ON at 4 °C. A freshly prepared polyacrylamide (PAA) solution (3% acrylamide (catalog no 161-0140, Bio Rad), 0.06 % bis-acrylamide (catalog no 161-0142 , Bio Rad), 0.05% ammonium persulphate (catalog no 161-0700, Biorad), 0.15 % TEMED, in PBS) containing 4% fluorescent beads (FluoSpheres, Invitrogen) was sandwiched between the silanized and the fibronectin-coated glass coverslip and polymerized at RT for 15 min, resulting in a 100 μm thick gel. The substrate was kept in PBS at 4 °C.

*Traction force microscopy*

15 kPa soft silicone substrates for TFM were prepared as described previously (3). In brief, CY52-276A and CY52-276B polydimethylsiloxane (Dow Corning, Toray) were mixed in a weight ratio of 1:1 (15 kPa) and then poured on glass bottom imaging dishes (Fluorodish, WPI) to obtain a layer of thickness around 100 µm. The gel covered dishes were spin-coated for 60 seconds at 400 rpm. The substrates were then cured at 80°C for 2 hours. Prior to seeding of the beads, the surface was silanized using a solution of 5% APTES ((3-Aminopropyl)triethoxysilane, Sigma-Aldrich) diluted at 10% in absolute ethanol for 10 min, then washed with absolute ethanol 3 times, before being dried at 80°C for about 10 min. 200 nm red carboxylated fluorescent beads (FluoSpheres, Invitrogen) were diluted within a 2:1000 ratio in water, and subjected to an ultrasonic bath for 10 min. The beads solution was then filtered using a 0.22 µm filter and incubated on the substrates for 15 min, protected from light. The dishes were finally washed with water 3 times and dried at 80°C for 3 minutes. Prior cell seeding, these substrates were coated with 50 µg/mL fibronectin or collagen (Sigma) for 45 min and washed 3 times with PBS. Adding Cy3-labelled fibronectin shows a uniform surface coating (images below).

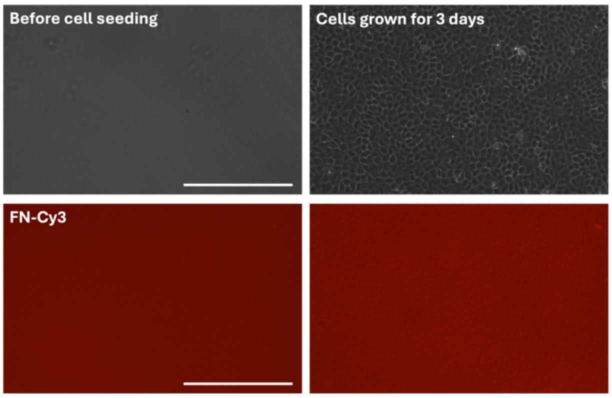


*Phase contrast and fluorescence images of 15 kPa PDMS coated with Cy3-labelled fibronectin. Left: Before cell seeding, right: after 3 days of cell culture. Scale bars 200 µm.*

*Micropatterning*

Polydimethylsiloxane (PDMS) stamps for micropatterning were prepared as described previously (*4*). Molds of the desired pattern were obtained using standard lithography methods. PDMS (Sylgard 184, Dow Corning) was prepared by mixing the base with a curing agent for a ratio of 1:10, poured over the mold, degassed, and then cured at 80°C for 2 h. Stamps were peeled of the mold and stored protected from light and humidity. Upon utilisation, the stamps surface was activated using plasma cleaning to make it hydrophilic, and a mixture of Cy3 conjugated fibronectin and regular fibronectin (50 µg/mL, Sigma) was then incubated covering the whole surface for 45 min, after which the surface was cleaned using a gentle air flow. The stamps were gently pressed against the bottom of PDMS covered culture dish for about 1 min for the pattern to imprint and with the stamps carefully lifted, the petri dishes were rinsed using PBS. The integrity of the patterns was verified using epifluorescence microscopy (Nikon). Patterns were then incubated with a solution of 2% Pluronic F127 (Sigma) for 1h to prevent cells from adhering on unstamped areas. The petri dishes were rinsed using PBS before seeding the cells.

<u>Cell culture</u>

The following cell lines were used:

- MDCK-II (ATCC CCL-34)
- MDCK-II LifeAct-Ruby (described in (*2*))
- MDCK-II E-cadherin knockout (clone B6P6, described in (*3*))
- MDCK-II E-cadherin knockout LifeAct-EGFP (clone B6P6, described in (*3*))
- MDCK-II E-cadherin/Cadherin 6 double knockout (clone D5, described in (*4*))
- MDCK-II E-cadherin overexpression (MDCK-E-cad GFP, described in (*5*)).
- MCF10A EGFP (described in (*6*))
- MCF10A E-cadherin knockout (described in (*7*))

MDCK-II cells were cultured in DMEM (GlutaMAX, high glucose and pyruvate, Life Technologies) supplemented with 10 % fetal bovine serum (FBS, Life Technologies) and 1 % penicillin-streptomycin (Life Technologies) at 37 °C with 5 % $CO_2$. MCF-10A cells were maintained in DMEM-F12 (#11039-021, Gibco) containing 10% penicillin-glutamine, 10 µg/mL human insulin (#I9278, Sigma Aldrich), 100 ng/mL cholera toxin (#C8052, Sigma-Aldrich), 0.5 mg/mL hydrocortisone (Sigma Aldrich), 5% horse serum, and 20 ng/ml EGF (PeproTeck) at 5% CO2 in an incubator at 37°C. Cells were passaged every 2-3 days using 0.05% Trypsin (catalog no 9002077, Merck). Before processing, the culture medium was aspirated and cells were rinsed with PBS to remove dead cells and debris.

<u>Sample preparation</u>

*Coculture experiments*

Competing cells were mixed in suspension using different ratios, seeded on a glass bottom imaging dish (Fluorodish) or a glass bottom imaging dish coated with 30 µg/ml Collagen G (Type 1, from calf skin, catalog no L7213, Sigma-Aldrich) or a PDMS or PAA substrate for traction force microscopy, and cultivated at standard conditions. After the desired cell density was reached, the culture medium was aspirated and cells were rinsed with warm PBS.

*Collision experiments*
5x10$^4$ cells were seeded in 80 μl culture medium into the wells of a 2-well culture inlet (catalog no 81176, Ibidi) on a glass bottom culture dish or a PDMS substrate for traction force microscopy. The cells were grown over night and the inlet was removed to allow the different populations to migrate towards each other.

*Ex-vivo culture of tumor patient derived xenografts*
Breast cancer patient derived xenografts were obtained from triple-negative breast tumors (HBCx-60 and BC953) and generated as previously described (*8*). After surgical excision of the tumor xenograft, tumoroids were isolated as previously described (*8*). Briefly, tissues were cut into small pieces and digested in RPMI1640 medium supplemented with collagenase 4mg/ml (Sigma-Aldrich) in 10 mM HEPES, 5% FBS, penicillin/streptomycin (1X), Glutamin (1X) for 1 hr at 37◦C on a rotating wheel at 150 rpm as previously described. Tumoroids were pelleted at 400g for 10 min. Then, tumoroids were incubated for 3-5 min at room temperature (RT) in DMEM/F12 / DNase (2U/μ l). Tumoroids were pelleted at 400g for 10 min. To remove fibroblasts, tumoroids were washed with DMEM/F12 medium and centrifugated at 400g for 3 seconds at RT until the supernatant is clear. Tumoroids were resuspended in DMEM with 10 mM HEPES, 5% FBS, 5μg/ml insulin, 10ng/ml Cholera toxin, hydrocortisone 1 mg/ml, penicillin/streptomycin (1X), Glutamin (1X). Around 800 tumoroids were plated on fluorodish coated with Fibronectin (50μg/ml) 37◦C. Metaplastic breast cancers contain epithelial (E-cadherin +, Vimentin -) and mesenchymal (E-Cadherin-, Vimentin +) cancer cells.

Stiffness measurements

Nanoindentation was used to measure the stiffness of cell monolayers. The indenter (Chiaro Nanoindenter, Optics 11 life, Netherlands) was connected to an epifluorescence microscope equipped with a 20x objective to visualize the indentation position. A soft probe with a small tip (k = 0.015 N/m, tip radius = 3 μm) was calibrated on glass before the measurement according to the manufacturers protocol.
Cell monolayers (pure MDCK WT, pure MDCK E-cad KO, coculture of WT and E-cad KO LifeAct-EGFP) were grown on glass and measured after confluency. Tumors were grown on fibronectin-coated glass or a thick layer of collagen. Before every measurement, the distance to the surface of the sample was determined automatically, and the probe was placed 5 μm above the surface. To measure the stiffness, the probe was pushed 1.5 μm into the sample for 4 sec and retracted afterwards. The matrix scan function was used with a typical stepsize of 25 μm and timelapse movies were acquired to trace back the indentation positions. For coculture measurements, E-cad KO LifeAct-GFP clusters were identified using the epifluorescence signal and the indentation positions were adjusted accordingly.
To determine the elastic modulus, the loading curve was analysed using the built-in software (DataViewer V2, Optics 11 life). The analysis is based on the Hertz model (Hertzian contact), which assumes a linear elastic response of the sample. The single fit method was used with a maximal load ($P_{max}$) of 90% and a Poisson's ratio of 0.5. Loading curves, which started in contact due to a failed determination of the surface were excluded automatically from the analysis.

Time-lapse microscopy

Confluent coculture monolayers or colliding cell populations were rinsed with warm PBS and fresh cultivation medium was added. For experiments where a pan-caspase inhibitor or a inhibitor of cell protrusions was used, 20 µm Z-VAD-FMK (catalog no tlrl-vad, InvivoGen) or 100 µM CK666 were added before starting the experiment. For experiments where actomyosin contractility was inhibited, 20 µM Blebbistatin (catalog no 203390, Sigma Aldrich) owere added to the cultivation medium during the experiment.
The dish was transferred to a live cell epifluorescence microscope (Biostation IM-Q, Nikon, equipped with a 10x or a 20x phase-contrast air objective and an incubation chamber) and incubated at 37 °C and 5% $CO_2$. The monolayer was imaged using phase contrast and the E-cad KO LifeAct-EGFP population and the fluorescent beads were imaged using epifluorescence. Time-lapse videos were taken at multiple positions every 15 minutes. In the case of traction force microscopy, cells were also removed at the end of the experiment by adding 200 µl of 10% SDS in the medium to obtain the relaxed state of the beads on the substrate.

Laser ablation experiments

MDCK WT and E-cad KO cells were seeded on glass-bottom imaging dish at a ratio of 50:50 and grown until reaching confluency so large islands of each cell type could be observed. Prior to wound induction, dishes were rinsed using warm PBS and provided with fresh culture medium. Laser ablation was done using a spinning disk CSU-X1 with a FRAP module (Yokogawa) equipped with a 40x/1.2 water-immersion objective. Briefly, holes the size of 3-4 cells were inducted within the area of the same cell type in the mixture, focusing a UV laser (355nm, 3-5ns pulse duration, laser power of 450nW) for 1s. Each samples were imaged during 15 s before the ablation and until 3 min after the ablation, using a 5 s intervals. The recoil velocity was measured by manually segmenting the edges holes over time and plotting the change in displacement of the edges of the ablated region.

Indirect immunostaining

Coculture or collision experiments which reached the desired cell density were rinsed with warm PBS. Fixation was carried out in 4% paraformaldehyde for 10 min at room temperature. Cells were permeabilized using 0.1% Triton X-100 in PBS for 5 min followed by 3 x 5 min washing in PBS. Samples were blocked with 1% BSA and 10 % FBS in PBS for 1 h at room temperature. Of note, the PDX tumoroids cells were permeabilized using 0.1% Triton X-100 in PBS for 10 min. All following primary antibodies were diluted 1:100 in blocking solution and incubated for 2 h at room temperature or over night at 4°C.

- anti E-cadherin mouse antibody (catalog no 610181, BD Biosciences) and
- anti E-cadherin clone ECCD2 for the 2D PDX staining (catalog no 1319000, ThermoFisher)
- anti α-catenin rabbit antibody (catalog no AB51032, Abcam)
- anti β-catenin rabbit antibody (catalog no 610156, BD Biosciences)
- anti paxillin rabbit antibody (catalog no AB32084, Abcam)
- anti phospho-myosin light chain 2 (pMLC2) rabbit antibody (catalog no 3671S, Cell Signaling)
- anti ZO1 rabbit antibody (catalog no 402300, Life Technologies)

- anti vimentin antibody (catalog no 8978, ThermoFisher)
- anti phospho-Histone H3 mouse antibody (Ser 10, catalog no 9706, Cell Signaling)
- anti desmoplakin mouse antibody (catalog no Cl.11-5F, Sigma)

The samples were washed 3 x 5 min with PBS and incubated with an anti rabbit (catalog no A31573, Life Technologies) or an anti mouse (catalog no A31571, Life Technologies) antibody conjugated to Alexa Fluor 647 diluted 1:200 in blocking solution for 2 h at room temperature. Subsequently, samples were washed 3 x 5 min with PBS. The actin cytoskeleton was visualized using Phalloidin-Alexa Fluor 568 (catalog no A12380, Life Technologies) diluted 1:200 in PBS and the nuclei using Hoechst 33342 (catalog no 62249, Thermo Fisher) diluted 1:2000 in PBS for 45 min at room temperature.

Confocal microscopy and data visualization

Fixed samples were mounted on a laser scanning confocal microscope (Zeiss LSM 980, Germany), equipped with a 63x oil objective and an Airyscan 2 module. MDCK WT cells expressing LifeAct-mCherry and E-cad KO cells expressing LifeAct-GFP were seeded on a glass-bottom imaging dish for live cell imaging. Timelapse videos were acquired using temperature (37 °C) and $CO_2$ control. Unless otherwise stated, all images or Z-stacks were acquired in Airyscan mode without further averaging, and an automated deconvolution was performed within the microscope software (ZEN blue). All images were visualized using Fiji (*9*) and brightness and contrast were adjusted. For Z-stacks, maximum intensity projections or side views were generated.

Image analysis

*Approximation of cell-cell adhesion strength*

Mixed cultures of MDCK WT and E-cad KO cells were stained for E-cadherin, α-catenin and β-catenin. To estimate the amount of recruited protein as an approximation of cell-cell adhesion strength, multiple line plots (length 10 µm, width 40 pixel) were acquired within the same image. The lines were placed manually perpendicular to and centred on junctions (WT-WT junction, E-cad KO-E-cad KO junction, WT-E-cad KO junction). The line plots were averaged and normalized to the highest average value.

*Quantification of focal adhesions*

Images of paxillin and LifeAct-EGFP (E-cad KO) were acquired at 4096 x 4096 pixel airyscan resolution and averaged four times. The LifeAct-EGFP signal was smoothed using first a 2 x 2 and then a 10 x 10 median filter. It was manually thresholder to generate a binary image of E-cad KO cells. A random forest classifier was trained using the pixel classification workflow in ilastik (*10*) to automatically segment focal adhesions based on the paxillin signal. Using the binary E-cad KO image, the focal adhesion segmentation was split into WT and E-cad KO, resulting in two separate binary images of WT and E-cad KO focal adhesions. In ImageJ, the 'Analyse Particles' function was used to quantify the focal adhesion area and to fit an ellipse to the focal adhesion segmentation. The major axis of the ellipse was used as a measure of the focal adhesion length.

*Quantification of phospho-myosin intensity*

Images of phospho-myosin and LifeAct-EGFP were acquired. Line plots spanning over 100 µm were acquired with the middle of the line placed perpendicularly on the cluster edge. These plots were normalized to the maximal intensity value and averaged.

*Quantification of cell height after tissue collision*
Large z-stacks of actin and LifeAct-EGFP spanning the interface and the bulk regions of both cell types were acquired. The interface was extracted based on LifeAct-EGFP signal. Side views of the actin channel were produced and binarized to obtain the cell height. The height was measured continuously over 400 µm with the tissue interface in the middle. Several positions were averaged.

*Cell segmentation and cell density quantification*
Large images of the coculture stained for ZO1, LifeAct-EGFP (E-cad KO), pHH3 and the nuclei were acquired. Nuclei were segmented using *StarDist* (*11*) at default settings and counted using Fiji (‘*Analyse Particles*’) to calculate cell densities. pHH3 positive cells were counted manually and their fraction was calculated based on the nuclei segmentation. Cells outlines were segmented based on the ZO1, Nuclei and LifeAct-EGFP signal using *Cellpose* (*12*).

*Interface convexity quantification*
Large island of e-cad KO cells within a confluent mixture of KO-WT cells had their interfaces segmented by hands using the LifeAct GFP channel, before and after adding blebbistatin. Convexity was calculated as the ratio between the perimeter of a given island and the perimeter of the corresponding convex shape (smallest polygon that can contain the shape of the island). Computation of the convex bounding region was done using MatLab image processing toolbox.

*Area fraction quantification*
The LifeAct-EGFP signal in time lapse movies of the confluent coculture or the collision experiment was converted to 8-bit grayscale and blurred using first a 2 x 2 median filter followed by a 5 x 5 median filter. A binary image was generated using manual thresholding. The binary image was blurred using a 5 x 5 median filter. The intensity of the whole image was measured at all timepoints. A one-hour rolling average was applied to compensate intensity fluctuations of the fluorescent lamp. The area was normalized to the starting value and plotted through time. The absolute area occupied by WT or E-cad KO cells was calculated by multiplying the fraction of E-cad KO cells (intensity divided by 255) with the area in the field of view (0,514188 $mm^2$ for the 10x phase contrast objective).

*Extrusion rate quantification*
Time lapse movies (phase contrast and LifeAct-EGFP signal of E-cad KO cells) of the coculture or the collision experiment were merged. A random forest classifier was trained by manual ground truth annotation using the pixel classification workflow in ilastik to automatically segment WT extrusions (= extrusion in phase contrast without LifeAct-EGFP signal) and E-cad KO extrusions (= extrusion in phase contrast with LifeAct-EGFP signal). Cell extrusions result in a strong increase in the phase contrast signal, with extruded cells appearing as bright spheres. The classifier was trained to detect these bright spheres and separate spheres close to each other, considering not only their brightness, but also their roundness and the smoothness of their edges. The rest of the image was annotated as background, in particular

cell divisions. When extruded cells die and fragment, they loose these features and were not considered any more (i.e. annotated as background). Cellular identities were attributed based on the LifeAct-EGFP signal. The classification resulted in a three-intensity image (WT extrusion, E-cad KO extrusion, background).
To assess the accuracy of the classifier, we calculated sensitivity (true positive rate) and specificity (true negative rate) for WT and E-cad KO extrusions:

| | **WT** | **E-cad KO** |
|---|---|---|
| *Sensitivity* | 100% | 92±7% |
| *Specificity* | 88±5% | 100% |

This means that WT cells are slightly oversegmented and very few E-cad KO cells are wrongly detected as WT, most likely due to inhomogeneities in LifeAct-EGFP expression strength. This means for the quantification of extrusion rates described below, that E-cad KO rates can be slightly under- and WT rates can be slightly overestimated. Due to decreasing performance with strongly increasing extrusion number (objects can not be separated any more), the analysis was limited to 24 hours.
The output was smoothed using two times a 2 x 2 median filter. In ImageJ, TrackMate (*13*) was used to detect and track extrusions, which were distinguished between WT and E-cad KO based on their intensity. A minimal area threshold (70 pixel) was set to exclude wrongly detected objects, too small for being a cell. To track extrusions through time and space, the simple LAP tracker was used with a gap closing distance of 40 pixel and a maximal linking distance of 40 pixel. The minimal length of a track was set to 3, corresponding to 45 min to exclude wrongly detected objects like floating debris or cell divisions. In the resulting output file, each track represents an extrusion event. The time stamp of the first spot defines the extrusion timepoint and its coordinates define the extrusion position.
To calculate the extrusion rate, the total number of WT or E-cad KO extrusions within two hour intervals were counted and divided by the area occupied by each cell type. Three consecutive time intervals were averaged, divided by two and normalized to achieve the number of extrusions per hour per $mm^2$.

*Spatial analysis of extrusion events*

Using the ImageJ plugin MorpholibJ (*13*) and binary images of the mixed culture, Euclidian distance maps were generated. In those distance maps, for a given position in the image, a pixel has a value equal to the distance to the closest interface between the two cells population. For each extrusion position, a distance from the interface was associated. Then using the random probability associated with any given distance as a normalization parameter, the probability distribution of being extruded knowing the distance from the interface was calculated for each cell type composing the mixture of the cocultures.

*Traction forces and stress measurements*

The bead images obtained during TFM manipulation were merged with corresponding reference bead images taken after SDS treatment. The resulting stack of images were pre-processed using the Image Stabilizer plug-in in ImageJ (*14*) and the illumination was corrected to remove background noise. Displacement field of beads was obtained using PIVlab (*15*), a particle image velocimetry toolbox developed in Matlab, with an interrogation window of 32x32 pixels and an overlap of 50%. Bead displacements were then correlated to a traction force field using Fourier transform traction cytometry (FTTC), known theoretical substrate stiffness, and a regularization parameter of 9x$10^{-9}$. From traction force field, we were able to infer stress tensor everywhere in the tissue using Bayesian inversion stress microscopy (*14*) with a regularization parameter $\Lambda = 10^{-6}$. Isotropic stress was calculated as half of the trace of the tress tensor. To generate heatmap of isotropic stress and traction force magnitude, a smoothing was applicated through linear interpolation.

*Stress inference from cell shape*
Results computed with BISM were also verified using the method based on cell shaped described in (*15*). Cell segmentation was done with cellpose on ZO-1 staining and cell identities were attributed according to E-cad KO lifeAct fluorescence.

*Local characterization of stress around extrusions and cell elimination stress maps generation*
Using a tuneable square interrogation window, the isotropic stress around each extrusion was extracted 2h before the extrusion until 1h after the extrusion. Extrusion positions were then filtered based on their distance from the interface, depending on the cell type, more or less than 30 µm. From all the remaining extrusion after the filtering, a median field of isotropic stress was computed, and from which, the mean stress evolution or the mean stress fluctuation evolution was plotted. From the same processed data were also generated the cell elimination stress maps for a moment in time between 40 and 30 minutes before the extrusion.

*Calculation of fluctuations*
Fluctuation of any parameter (isotropic stress, traction force, or bead displacement) was characterized using the susceptibility χ as described in (*16*) and used in other experimental analysis (*17*). Briefly, for a given physical parameter A distributed inside a population of N cells, the susceptibility can be computed as $\chi_A = Var(A) * N$. In some case where the number of cells was not convenient to access, the number of pixel in the considered area was using as a proxy to compute the susceptibility, given that all the cells were sharing the same average area.

### Statistical analysis
All plots/graphs show the mean. All error bars show the standard deviation. All statistical tests were performed using GraphPad Prism (Version 9.5.0).

### Computational model
#### 3D phase-field model

We use a recently developed three-dimensional (3D) phase-field model for active cell layers (*18*). Within this framework cells are represented as 3D deformable particles which dynamically adapt their shape in response to active stresses as well as interactions forces with other cells and the underlying substrate. In this vein, we consider a cellular monolayer consisting of $N_{cell}$ cells on a rigid substrate with its surface normal $\vec{e}_n(= \vec{e}_z) = \vec{e}_x \times \vec{e}_y$ and periodic boundaries in both $\vec{e}_x$ and $\vec{e}_y$, where $(\vec{e}_x, \vec{e}_y, \vec{e}_z)$ constitute a global orthonormal basis. Each cell $i$ is represented by a three-dimensional (3D) phase-field, $\phi_i = \phi_i(\vec{x}, t)$ and initialized with radius $R_0$. The dynamics associated with relaxation of cell interface follows a time-dependent Ginzburg-Landau model with an extra advective term:

$$\partial_t\phi_i + \vec{v}_\mathrm{i}\cdot\vec{\nabla}\phi_i = -\Gamma\frac{\delta F}{\delta\phi_i},\ i\ =\ 1,\dots,N_{cell}\ [1]$$

Where $\Gamma$ is the mobility coefficient. Furthermore, the advective term $\vec{v}_i\cdot\vec{\nabla}\phi_i$ updates the location of $\phi_i = \phi_i(\vec{x},t)$ for each time step and each cell $i$ with velocity $\vec{v}_i$. The free energy functional reads (*20*):

$$F\ =\ \sum_i^{N_{cell}}\frac{\gamma^i}{\lambda}\int d\vec{x}\left\{4\phi_i^2(1-\phi_i)^2+\lambda^2\left(\vec{\nabla}\phi_i\right)^2\right\}$$

$$+\sum_i^{N_{cell}}\mu\left(1-\frac{1}{V_0}\int d\vec{x}\,\phi_i^2\right)^2+\ \sum_i^{N_{cell}}\sum_{j\neq i}\frac{\kappa_{cc}}{\lambda^2}\int d\vec{x}\,\phi_i^2\phi_j^2$$

$$+\sum_i^{N_{cell}}\sum_{j\neq i}\omega_{cc}^i\int d\vec{x}\left(\vec{\nabla}\phi_i\cdot\vec{\nabla}\phi_j\right)++\ \sum_i^{N_{cell}}\frac{\kappa_{cs}}{\lambda^2}\int d\vec{x}\,\phi_i^2\phi_w^2$$

$$+\sum_i^{N}\omega_{cs}^i\int d\vec{x}\left(\vec{\nabla}\phi_i\cdot\vec{\nabla}\phi_w\right)\ [2]$$

As such the free energy stabilizes cell interface and includes mechanical properties of the cells such as cell cortex tension ($\gamma^i$), as well as gradient contributions ( $\vec{\nabla}\phi_i$) that account for, and distinguish between, cell-cell ($\omega_{cc}^i$) and cell-substrate ($\omega_{cs}^i$) adhesions. In addition to the cortex tension and adhesion terms, compressibility ($\mu$), puts a soft constraint on the cell around $V_0 = (4/3)\pi R_0^3$ and $\kappa$ captures repulsion between cell-cell (subscript cc) and cell-substrate (subscript cs) and $\phi_w$ denotes a static phase-field representing the substrate (see Fig. SX for a schematic). Based on this free energy functional, the interior and exterior of cell i corresponds to $\phi_i = 1$ and $\phi_i = 0$ respectively, connected by a diffuse interface parameterized by a length $\lambda$. To resolve the forces generated at the cellular interfaces, we utilize an overdamped dynamics:

$$\vec{T_i} = \xi\vec{v_i} - \vec{F}_i^{\mathrm{sp}} = -\int d\vec{x}\left(\mathbf{\Pi}^{int}\cdot\vec{\nabla}\phi_i\right)\ [3]$$

where $\vec{T_i}$ denotes traction as defined for Bayesian Inversion Stress Microscopy in (*14*), $\xi$ is substrate friction and $\vec{F}_i^{\mathrm{sp}} = \alpha\vec{p_i}$ represents self-propulsion forces due to polarity, constantly pushing the system out-of-equilibrium. In this vein, $\alpha$ characterizes the strength of polarity force and

$$\mathbf{\Pi}^{int} = \left(\sum_i^{N_{cell}}-\left(\frac{\delta F}{\delta\phi_i}\right)\right)\mathbf{1} + \left(\sum_i^{N_{cell}}-\left(\zeta_S^i\phi_i\boldsymbol{S}_i\right)\right) + \left(\sum_i^{N_{cell}}-\left(\zeta_Q^i\phi_w\boldsymbol{Q}_w\right)\right)\ [4]$$

where $\boldsymbol{S}_i = -\int d\vec{x}\left\{\left(\vec{\nabla}\phi_i\right)^T\vec{\nabla}\phi_i\right\} + \left(\frac{1}{3}\right)Tr\int d\vec{x}\left\{\left(\vec{\nabla}\phi_i\right)^T\vec{\nabla}\phi_i\right\}$ and $\boldsymbol{Q}_w = \left(\frac{d}{d-1}\right)\left(\vec{n}\otimes\vec{n} - \frac{1}{d}(\vec{n})^2\mathbf{1}\right)$ where $d = 3$ is the dimension, $\zeta_Q^i$ and $\zeta_S^i$ are the strength of cell-substrate and cell-cell active stresses, respectively, and $\boldsymbol{I}$ is the identity tensor. In the model, $\vec{n}$ represent the orientation of stress fibers of the substrate, e.g. $\vec{n} = (\cos\upsilon, \sin\upsilon, 0)$ and $\upsilon = 0$ on the surface of the substrate. Furthermore, the dynamics of cell polarity is introduced based on contact inhibition of locomotion (CIL) (*19*, *20*) by aligning the polarity of the cell to the direction of the total interaction force acting on the cell (24). As such, the polarization dynamics is given by:

$$\partial_t\theta_i = -\frac{1}{\tau_{pol}}\Delta\Theta_i + D_r\eta(t)$$

where $\theta_i \in [-\pi, \pi]$ is the angle associated with polarity vector, $\vec{p_i} = (\cos\theta_i, \sin\theta_i, 0)$ and $\eta(t)$ is a standard Gaussian white noise with zero mean unit variance, $D_r$ is rotational diffusivity, $\Delta\Theta_i$ is the angle between $\vec{p_i}$ and $\vec{T_i}$, and positive constant $\tau_{pol}$ sets the alignment time scale. Finally, we compute a coarse-grained stress field $\boldsymbol{\sigma}^i = \boldsymbol{\sigma}^i(\vec{x}, t)$ that encodes both active and passive contributions on a discretized domain, for node $i$:

$$\boldsymbol{\sigma}^i = \frac{1}{a_0^3} \sum_j^{N_i} \vec{r}^{ij} \otimes \vec{T}^j$$

where $a_0 = 1$ is the grid size and correspond to spatial domain discretization and $\vec{r}^{ij} = (\vec{x}^i - \vec{x}^j)$ and $N_i$ is the number of nearest neighbors at node i. A negative stress value indicates compression and a positive value, tension.

3D phase-field model simulation details

Specifically, we consider a cellular monolayer consisting of $N = 400$ cells on a rigid substrate. Cells are initiated on a two-dimensional simple cubic lattice and inside a cuboid of size $L_x = L_y = 320, L_z = 64$ and radius $R_0 = 8$. The total number of time steps in the simulations are $n_{sim} = 15{,}000$.

Unless specified otherwise, time $\tilde{t} = t/\tau$ where $\tau = (2R_0)/\overline{v}$ and $\overline{v}(= 0.02)$ in simulation units is the average speed of cells and $\tau$ represents the characteristic time for a cell to move a distance equivalent to its size. With this normalization and for a typical MDCK cell speed of $\sim 20\ \frac{\mu m}{h}$ and cell size $\sim 20\mu m$. The physical properties are as follows unless specified otherwise: $\gamma = 0.008$, $\mu = 45$, $\xi = 1$, $\zeta_S = +4E-5$ and $\zeta_Q = -0.01$.

Energetics Model

The energetics model is based on the work done to extrude the cell. There are two contributions to the work: adhesion and surface tension. The work needed to increase the surface area by $dA$ against surface tension is given by $kdA_{tot}$. In contrast, for adhesion, the cell is doing work to increase the surface area. The total work is therefore:

$$dW = -\omega_i dA_i + kdA_{tot} \text{ [6]}$$

The subscript '$i$' refers to the cell-cell adhesion and the cell-substrate adhesion. The negative sign in front of '$\omega_i$' indicates that the cell is doing the adhesive work. Consider the various contributions to the adhesion energy for cell type-I. There is the cell-substrate adhesion energy of cell type-I and of cell type-II, cell-cell adhesion energy of cell type-I with both a type-I and type-2 neighbor. Each energy is proportional to the change in the respective contact area. Expanding,

$$dW^I = -\omega_s^I dA_s^I - \omega_c^I dA_{cc}^I - \omega_c^{I-II} dA_{cc}^{I-II} + \omega_s^{neigh} dA_s^{neigh} + k^I dA_{tot}^I \text{ [6]}$$

The 1st term represents the cell-substrate adhesion of the cell at the interface. The second term represents the cell-cell adhesion of the cell with neighbors of the same type and the third term is cell-cell adhesion with a neighbor of the other type. When the cell is extruded, the substrate area it used to occupy is occupied by its neighbors. Therefore, there is an additional energy, the 4th term, from the substrate energy of the neighbors. The last term is the intrinsic stiffness of the cell.

Half the cell is assumed to be in contact with the cell of the same type and half with the other type. A factor of 0.5 is chosen for simplicity. We assume the winning cell pushes the losing cell from the bottom, and hence occupies the entire substrate contact area. The energy difference $dW^{II} - dW^{I}$ decides which of the cell types wins at an interface. Notice that the $\omega_c^{I-II}$ term will be the same in both the work function $dW^{I}$ and $dW^{II}$. It will therefore be dropped. The work done is now:

$$dW^{I} = (\omega_s^{II} - \omega_s^{I})dA_s^{I} - \omega_c^{I}dA_{cc}^{I} + k^{I}dA_{tot}^{I} [6]$$

The change in area is directly calculated using the surface areas of the shapes shown in schematic (Fig. S13), in which each shape is assumed to have the same volume. The cell is initially assumed to be a cylinder, whose basal radius is allowed to vary. Stress fluctuations lead to the shrinking of the basal radius, leading to a cone. The work needed to break the cell-cell adhesion bond is calculated by changing the shape of the cell from a cone to a sphere, again with constant volume. The radius of the sphere and the apical radius is both assumed to be constant at 10 microns. The volume is therefore $\frac{4}{3}\pi 10^3 \sim 4188.8\ \mu m^3$.

The difference in work in Figure 2H of the main text is normalized by the quantity, $dW_0^k$. This is the energy released by a cell to go through the process if there was no cell-cell or cell-substrate adhesion.

4. A. Glentis, C. Blanch-Mercader, L. Balasubramaniam, T. B. Saw, J. d'Alessandro, S. Janel, A. Douanier, B. Delaval, F. Lafont, C. T. Lim, D. Delacour, J. Prost, W. Xi, B. Ladoux, The emergence of spontaneous coordinated epithelial rotation on cylindrical curved surfaces. *Science Advances* **8**, eabn5406.

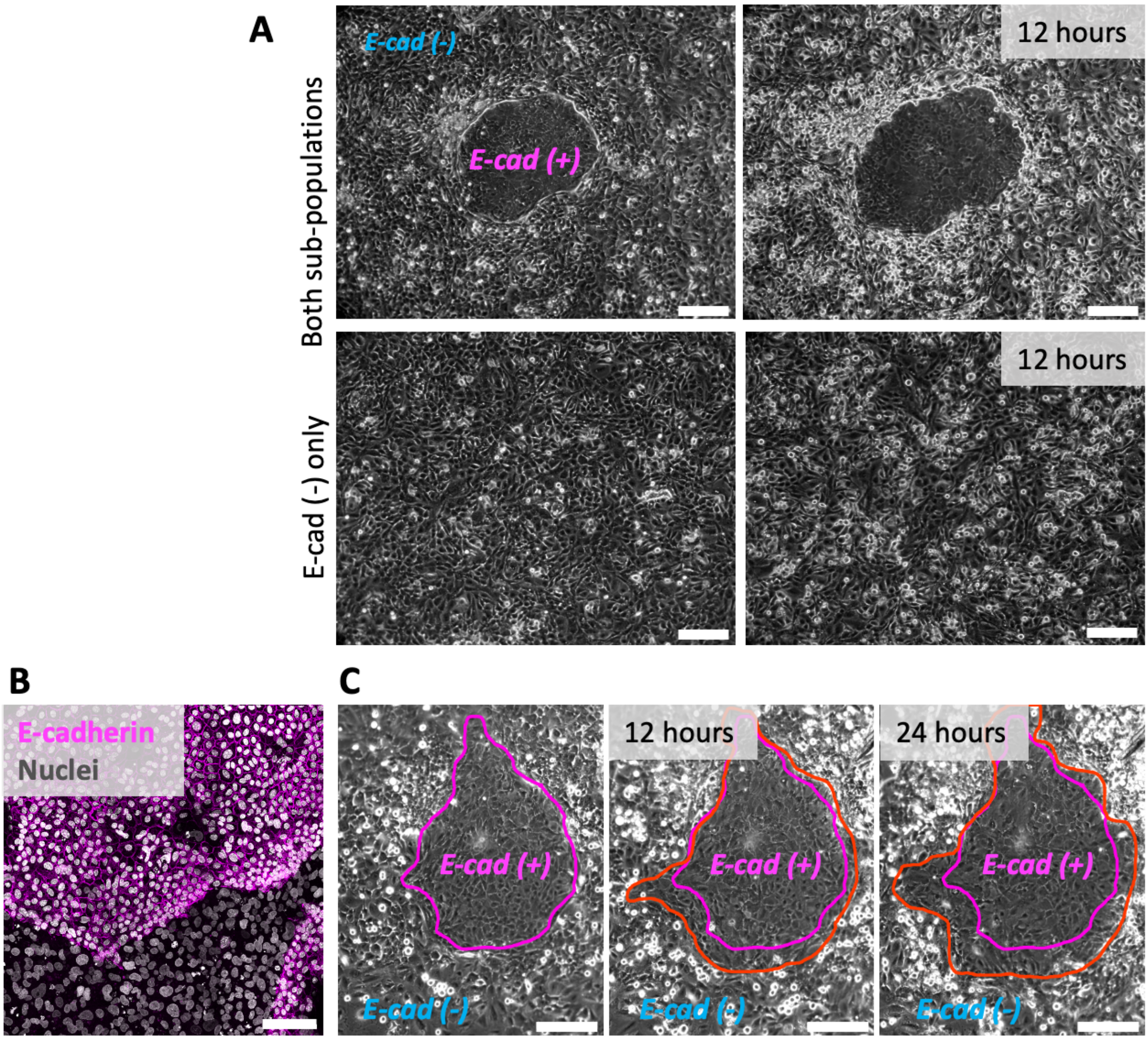


**Fig. S1: E-cad⁻ triple negative tumor cells are viable and sample from second patient.** (**A**) Upper: Mix of both sub-populations. The E-cad$^+$ population is spreading, leading to eliminations of E-cad$^-$ cells in its vicinity. Lower: E-cad$^-$ cells grow to high densities independent of interactions with E-cad$^+$ cells. Over time, elimination at random positions can be observed, but fewer than when interacting with the E-cad$^+$ sub-population. (**B**) Confocal image of metaplastic breast cancer cells derived from a second patient stained for E-cadherin (magenta). E-cad$^+$ and E-cad$^-$ cells have sorted. (**C**) Phase contrast images of cells corresponding to (**B**). Magenta line shows initial boundaries of the sub-populations red lines show boundaries at the given timepoint. Representative images from N=3 independent experiments. Scale bars 200 µm.

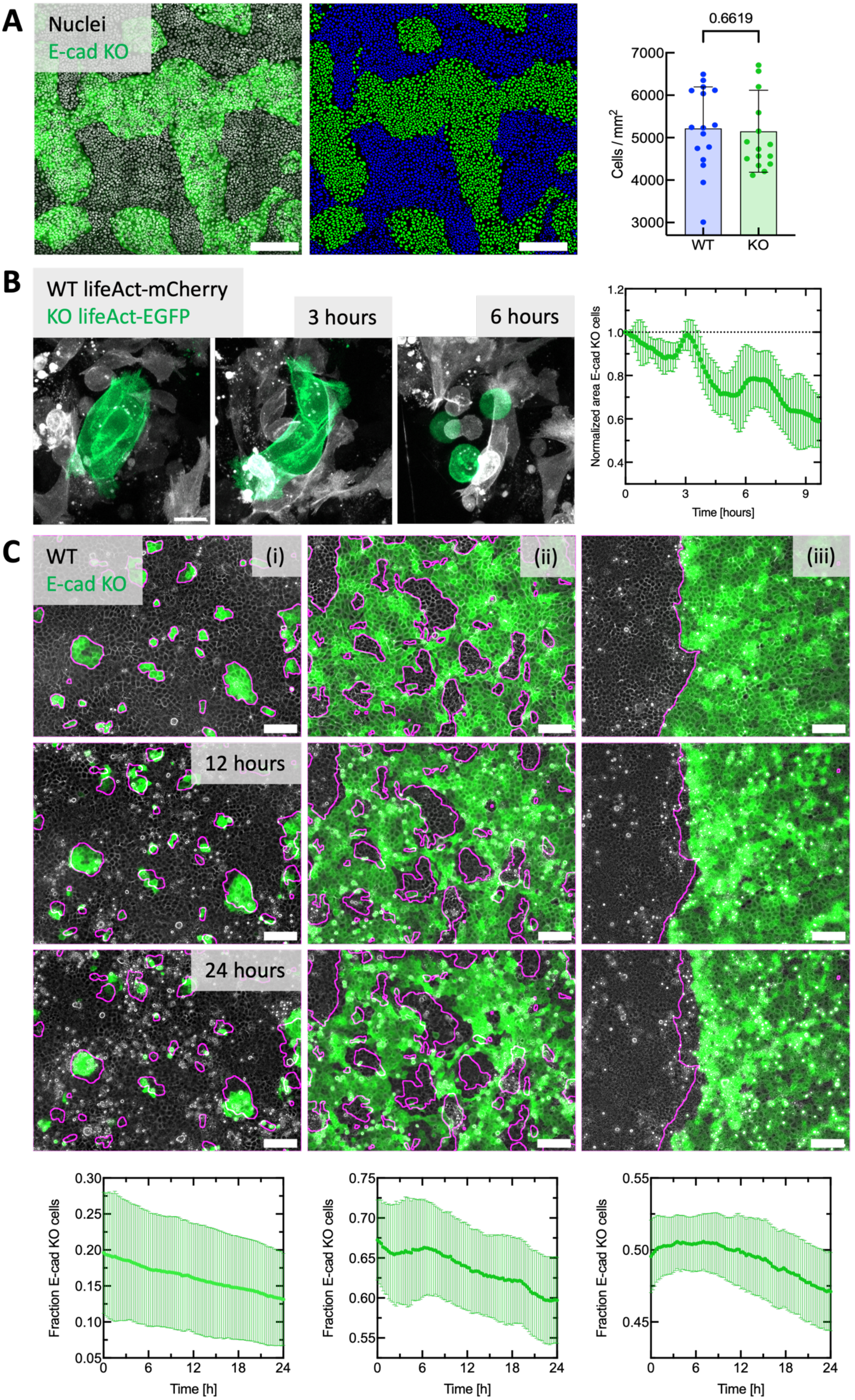
A
Nuclei
E-cad KO
0.6619
Cells / mm$^2$
7000
6000
5000
4000
3000
WT
KO
B
WT lifeAct-mCherry
KO lifeAct-EGFP
3 hours
6 hours
Normalized area E-cad KO cells
1.2
1.0
0.8
0.6
0.4
0
3
6
9
Time [hours]
C
WT
E-cad KO
(i)
(ii)
(iii)
12 hours
24 hours
Fraction E-cad KO cells
0.30
0.25
0.20
0.15
0.10
0.05
0.75
0.70
0.65
0.60
0.55
0.50
0.45
0
6
12
18
24
Time [h]

**Fig. S2: E-cad KO cells lose independent of ratio or level of sorting.** (**A**) Confocal image of nuclei (white) and E-cad KO cells (green). Middle: Corresponding nuclei segmentation using StarDist. The cellular identity was determined based on the E-cad KO fluorescence. Right: Average cell densities in mixed cultures. Each datapoint shows the fraction for one FOV. n=16 FOV from N=2 independent experiments. P-value from unpaired t-test. (**B**) Left: Snapshots from Airyscan confocal movies of WT cells expressing LifeAct-mCherry and E-cad KO cells expressing LifeAct-EGFP. The E-cad KO cells are eliminated by LifeAct expressing WT cells. Right: Area quantifications corresponding to images on the left from n=4 movies from N=1 experiment. (**C**) Brightfield and fluorescence images of WT and E-cad KO cells (green) at the beginning of the experiment and 12 and 24 hours later. Magenta lines show initial cluster sizes to visualize cluster progression over time. Bottom row: Area development of E-cad KO cells. (**i**) E-cad KO in minority. (**ii**) E-cad KO in majority. (**iii**) Collision of E-cad KO and WT tissues. Movies are analysed shortly after contact formation. Area quantifications from (**i**) n=8 movies from N=3 independent experiments, (**ii**) n=6 movies from N=3 independent experiments, (**iii**) n=4 movies from N=2 independent experiments. Error bars show the standard deviation. Scale bars 200 µm (A), 100 µm (C), 10 µm (B).

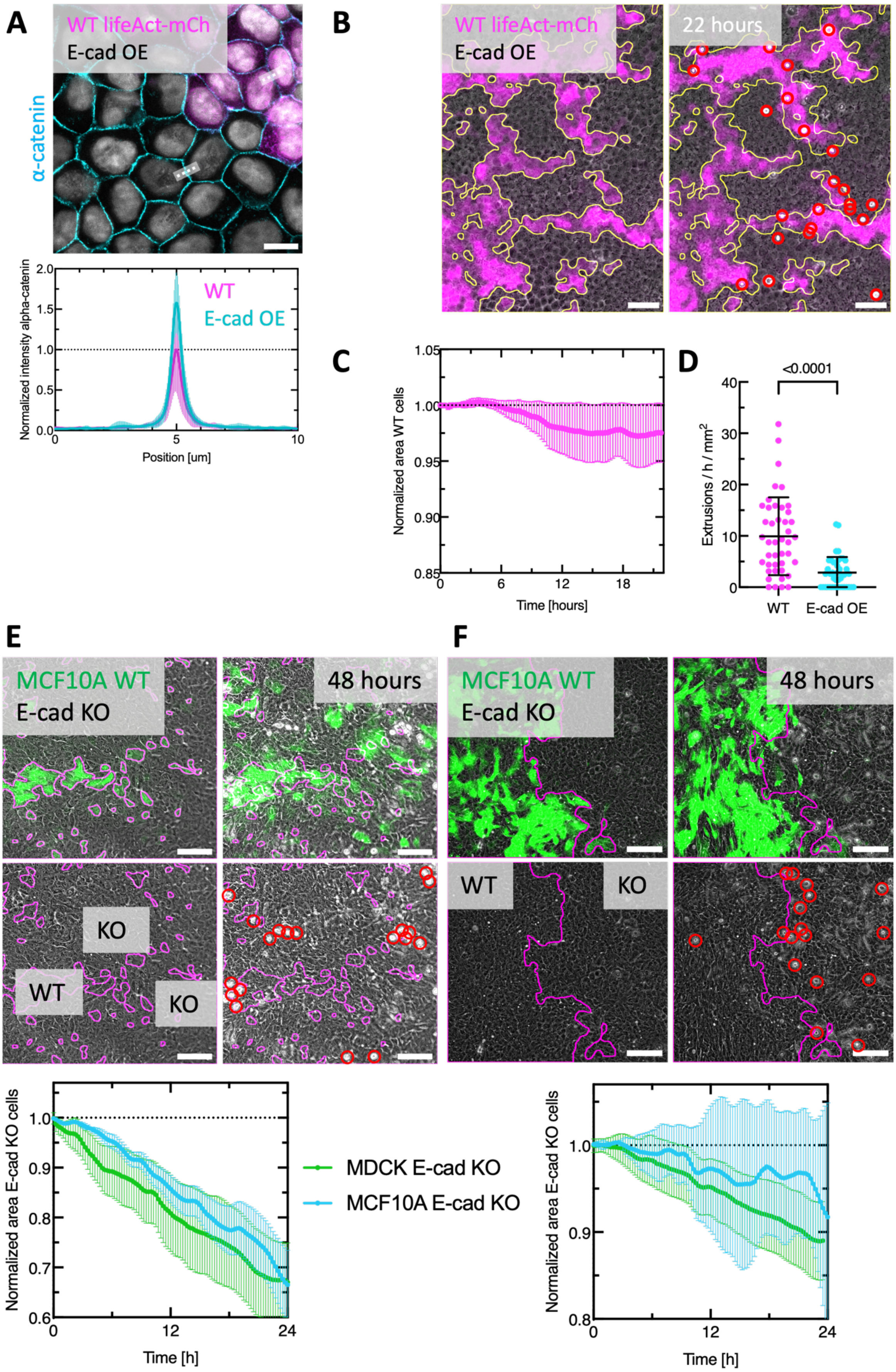
A
WT lifeAct-mCh
E-cad OE
α-catenin
Normalized intensity alpha-catenin
WT
E-cad OE
Position [um]
B
WT lifeAct-mCh
E-cad OE
22 hours
C
Normalized area WT cells
Time [hours]
D
<0.0001
Extrusions / h / mm²
WT
E-cad OE
E
MCF10A WT
E-cad KO
48 hours
KO
WT
KO
F
MCF10A WT
E-cad KO
48 hours
WT
KO
Normalized area E-cad KO cells
Time [h]
MDCK E-cad KO
MCF10A E-cad KO
Normalized area E-cad KO cells
Time [h]

**Fig. S3: MDCK E-cad-overexpressing cells eliminate MDCK WT cells and MCF10-A E-cad KO cells get eliminated by WT cells.** (**A**) Increased alpha-catenin intensity at junctions between E-cad-overexpressing (E-cad OE) cells (described in *Adams et al., JCB, 1998*) compared to WT cells. Upper: representative confocal image. Lower: Junctional alpha-catenin intensity normalized to WT cells. (**B**) Representative images of mixed culture of MDCK WT lifeAct-mCherry and E-cad OE cells. Yellow outlines show initial clusters, red circles mark extrusions. (**C**) Area development of MDCK WT lifeAct-mCherry cells in competition with E-cad OE cells. n=6 movies from N=2 independent experiments (**D**) Extrusion rates of WT lifeAct-mCherry and E-cad OE cells. Each datapoint shows one time interval from n=4 movies and N=2 independent experiments. P-value from unpaired t-test. (**E**) Mixed culture and (**F**) collision of MCF10-A EGFP (green) and MCF10-A E-cad KO (not fluorescent) cells. Magenta outlines show initial clusters. Red circles mark extrusions. Below: Area development of MCF10A E-cad KO cells compared to MDCK E-cad KO cells (shown in **Figure 1**) normalized to initial value. MCF10A data: n=5 movies from N=2 independent experiments for coculture and collision. Scale bars 100 µm (C,D); 50 µm (B); 10 µm (A).

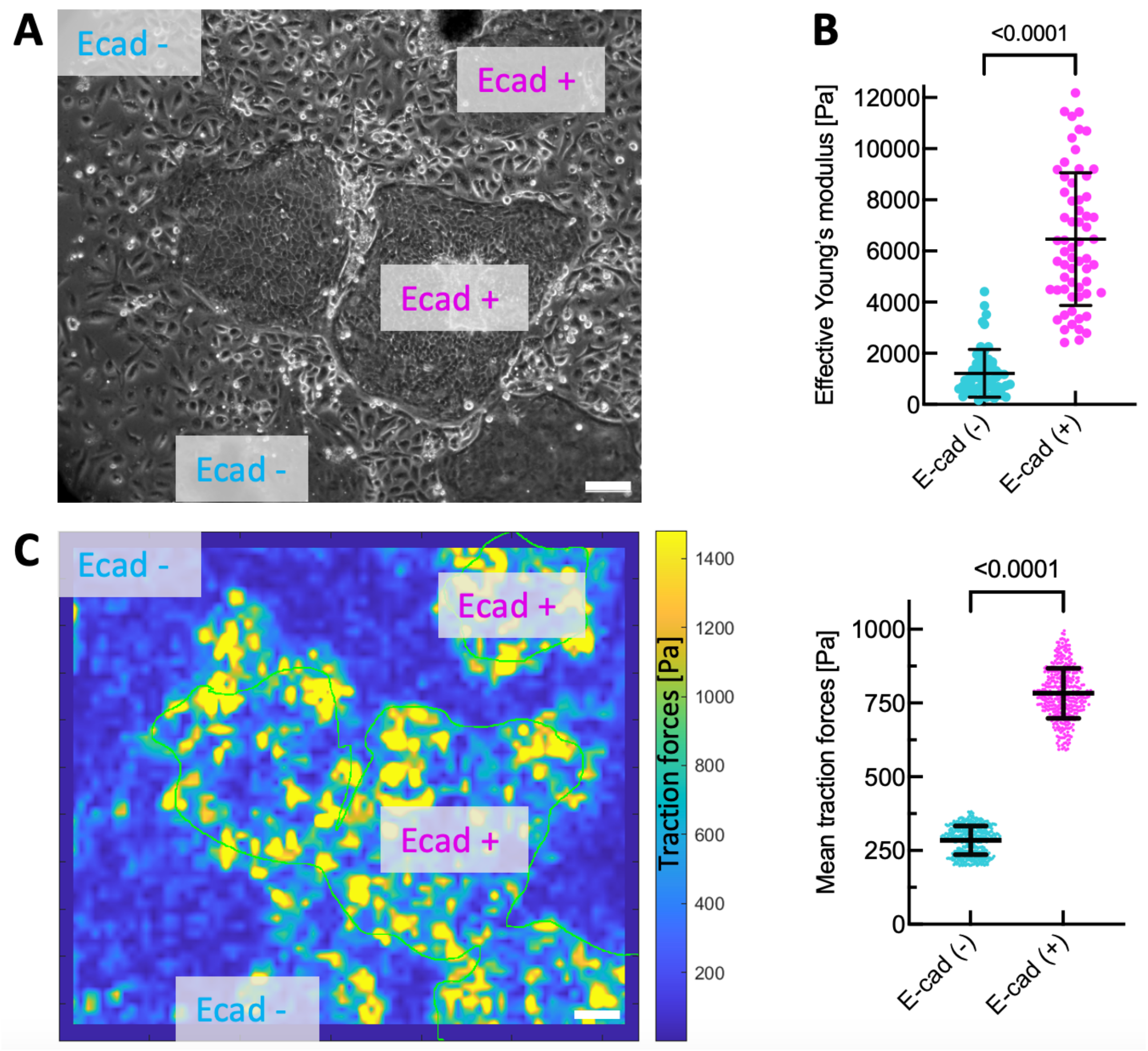


**Fig. S4: Mechanics of intratumoral cell competition.** (**A**) Brightfield image of E-cad (-) and E-cad (+) tumor clusters, as shown in **Figure 1**. (**B**) Cell stiffnesses measured with nanoindentation. Each datapoint shows one measurement. N=2 independent experiments and n=57 indentations. (**C**) Map of traction forces corresponding to (**A**). Green lines show cluster outlines. Right: Average traction forces. Each datapoint shows the average value from one field-of-view at one timepoint. N=2 independent experiments and n=408 timepoints. Scale bars 100 µm.

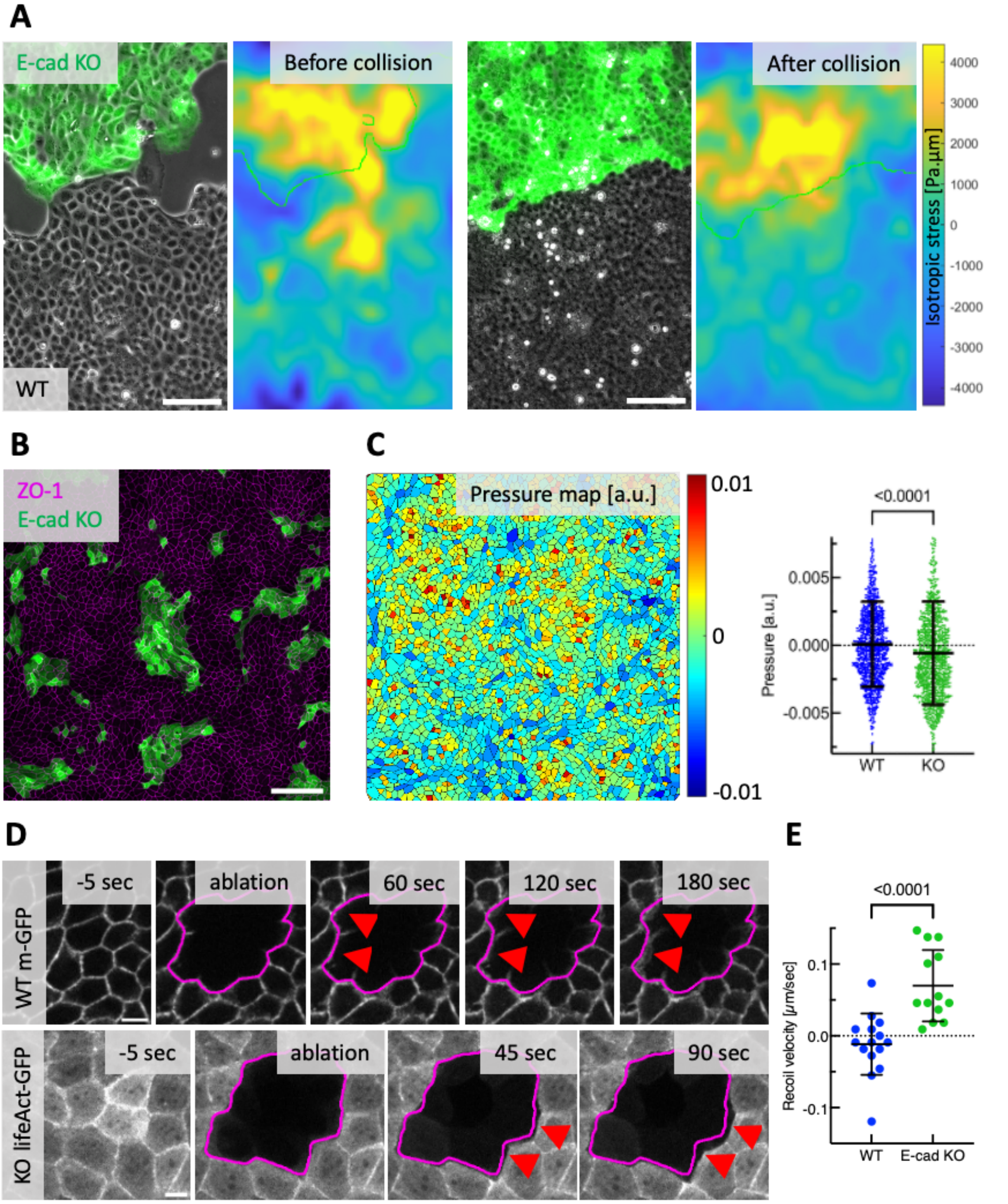


**Fig. S5: Different verifications of stress measurements.** (**A**) Isotropic stresses before and after collision of E-cad KO (green) and WT tissues. Before collision, tension is highest at the migration front of both cell types. After collision, the WT cells are under compression. (**B**) Example confocal image of mixed culture (LifeAct-E-cad KO in green) stained for cell boundaries (ZO-1, magenta). (**C**) Example image and average relative pressure inferred based on cell shape, as described in Kong et al., Sci. Rep. 2019. Each datapoint represents a single cell. n=1000 cells representing N=2 independent experiments. (**D**) Representative images of WT cells expressing CAAX-GFP mixed with E-cad KO cells expressing lifeAct-GFP before and after laser ablation. (**E**) Comparison of the recoil velocity measured after the ablation. Each datapoint represents one ablation representing N=2 independent experiments. P-value from unpaired t-test. Scale bars 100 µm (A,B); 10 µm (D).

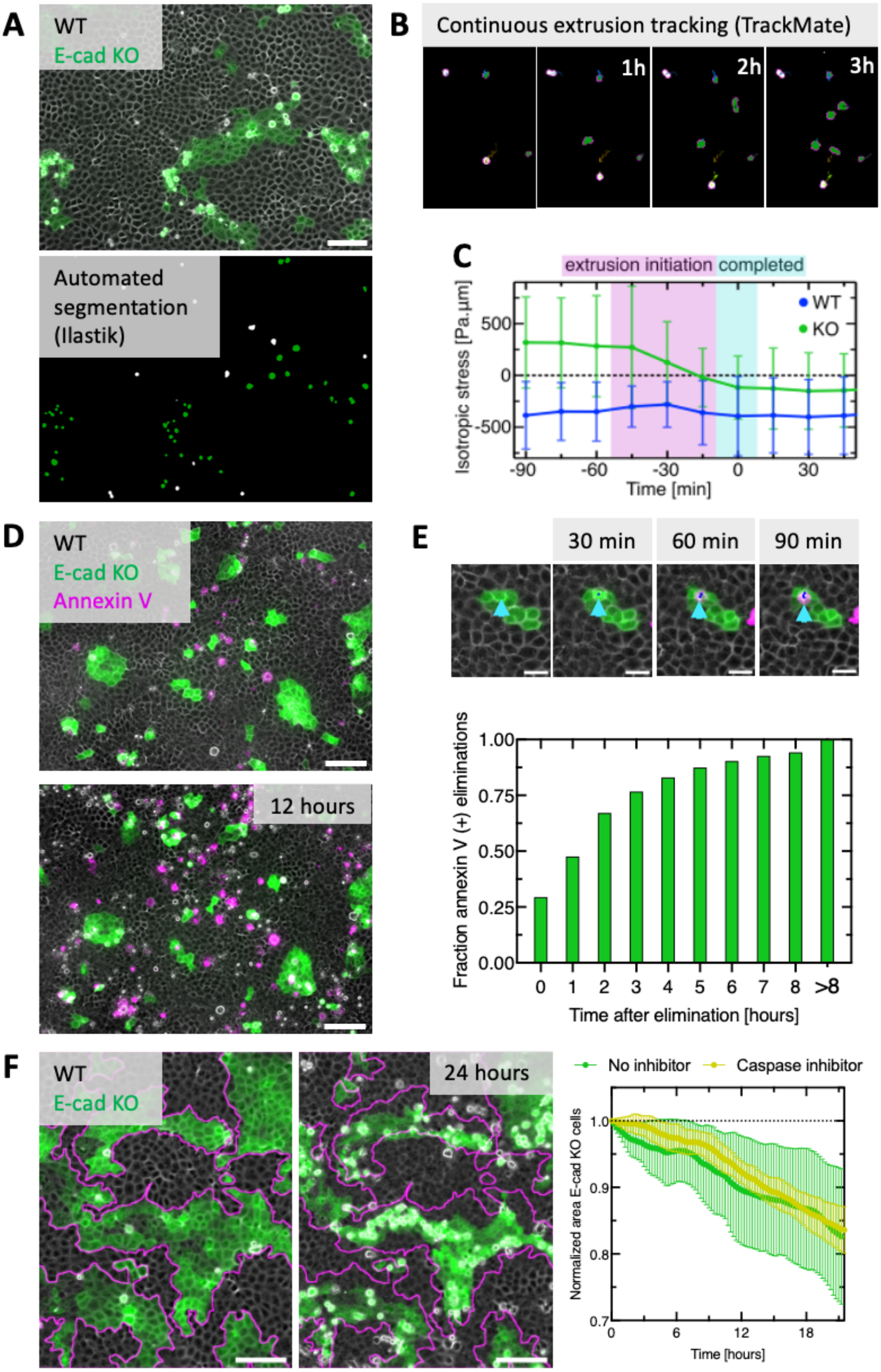
A
WT
E-cad KO
Automated segmentation (Ilastik)
B
Continuous extrusion tracking (TrackMate)
1h
2h
3h
C
extrusion initiation
completed
WT
KO
Isotropic stress [Pa.µm]
500
0
-500
-90
-60
-30
0
30
Time [min]
D
WT
E-cad KO
Annexin V
12 hours
E
30 min
60 min
90 min
Fraction annexin V (+) eliminations
1.00
0.75
0.50
0.25
0.00
0
1
2
3
4
5
6
7
8
>8
Time after elimination [hours]
F
WT
E-cad KO
24 hours
No inhibitor
Caspase inhibitor
Normalized area E-cad KO cells
1.0
0.9
0.8
0.7
0
6
12
18
Time [hours]

**Fig. S6: Cell elimination is independent of compressive stress or cell death.** To quantify E-cad KO and WT cell extrusions and determine their position, we developed an automated workflow: (**A**) Exemplary raw data (merged brightfield and fluorescent signal) and segmentation of extrusions (white = WT, green = E-cad KO). The supervised learning algorithm ilastik was trained to detect and classify extrusions based on both signals. Details about the training and the accuracy can be found in the Methods section. (**B**) To determine accurate extrusion positions and avoid counting extrusions multiple times, the extrusions were tracked through time using TrackMate. (**C**) Average isotropic stress before E-cad KO and WT extrusions. T=0 indicates detection, i.e. completion of the extrusion process. Stresses were measured within a square of size 60 µm around one extrusion event for different time points. n=726 (E-cad KO) and n=334 (WT) extrusions in n=7 movies from N=2 independent experiments. Error bar shows s.d. (**D**) Cell death is detected by the annexin V singal (magenta) in mixed culture. (**E**) Quantification of cell fate. For each extrusion, we calculated the time from cell elimination until detection of the annexin V signal. The plot shows the fraction annexin V-positive cells after the time of extrusion. n=11327 extrusions from n=17 movies and N=4 independent experiments. (**F**) Representative images of mixed culture treated with a pan-caspase inhibitor (Z-VAD-FMK, 20 µM). Magenta line shows initial cluster boundaries. Right: Area quantification compared of the standard experimental condition. N=1 caspase inhibitor experiment and n=5 positions. Scale bars 100 µm (A, D, F); 25 µm (E).

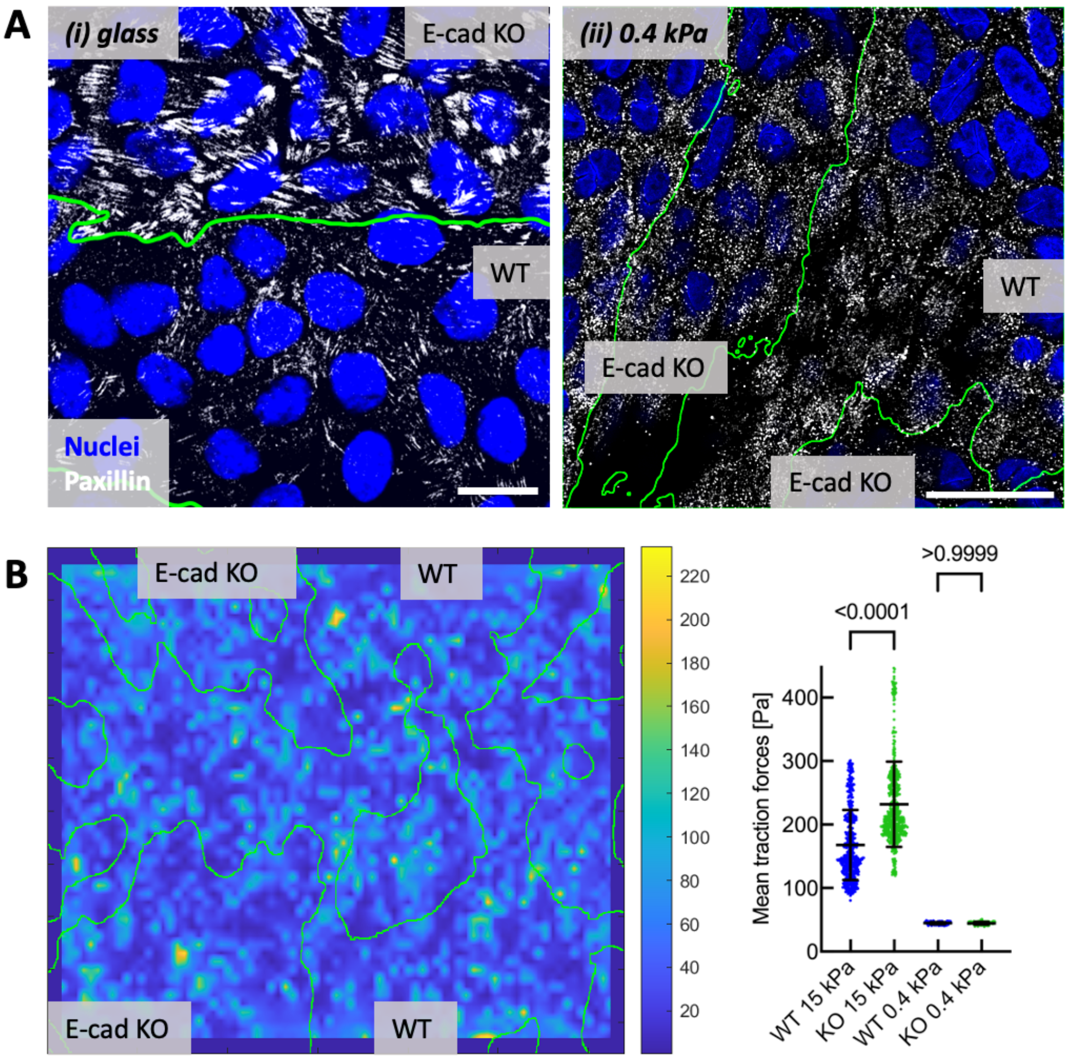


**Fig. S7: Mixed culture with low cell-substrate adhesion.** (**A**) Confocal images of focal adhesions (Paxillin, white) in mixed culture grown on (i) glass or on (ii) soft substrate (370 Pa, PAA). (**B**) Left: map of traction forces on soft substrate. Right: Average traction forces on different substrate stiffnesses. Green lines show outlines of E-cad KO cluster. Each datapoint represents one average map. n=568 positions from N=4 independent experiments (15 kPa) and n=170, N=2 (370 Pa). P-values from Kruskal-Wallis test corrected for multiple comparisons (Dunn's test). Scale bars 20 µm.

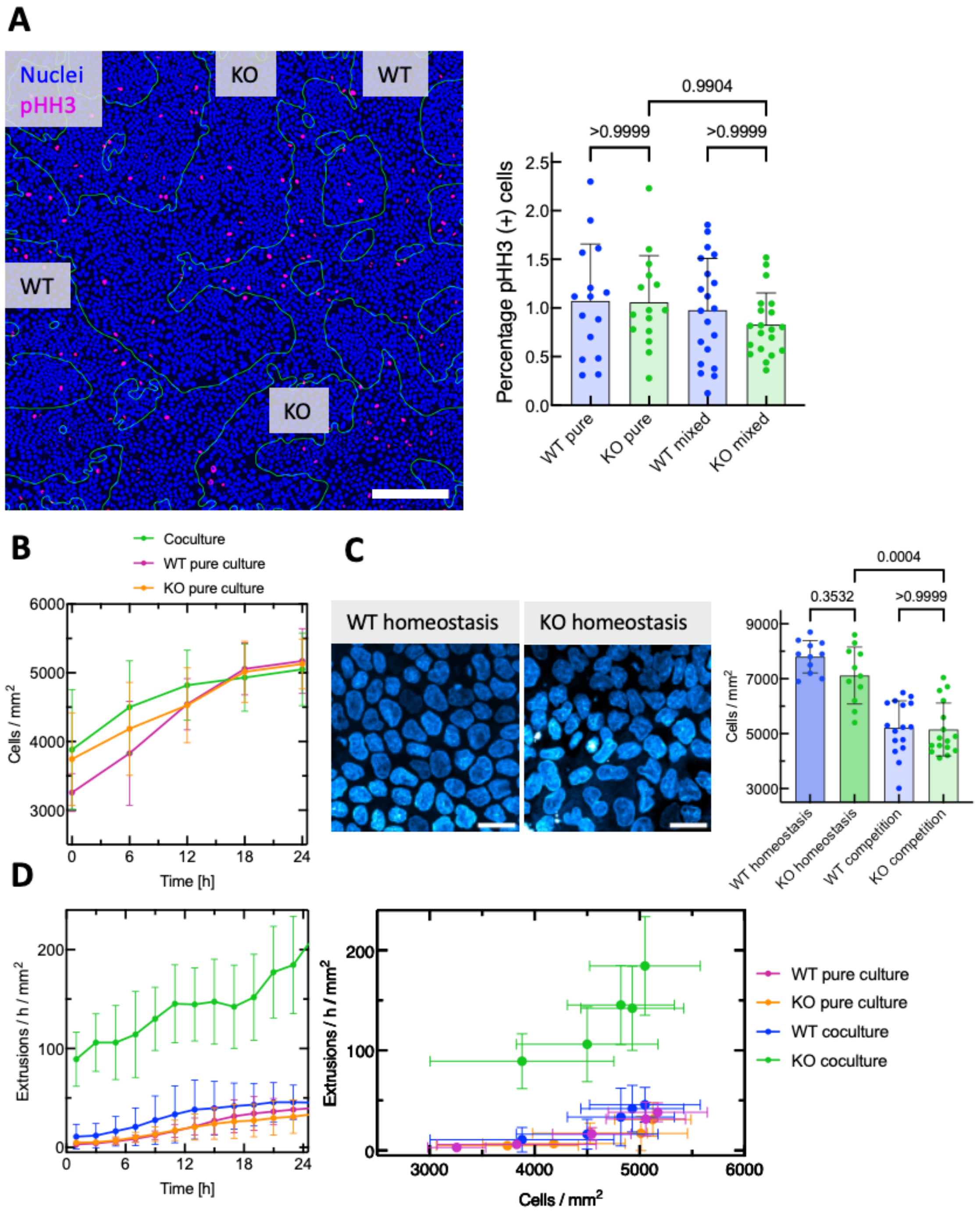


**Fig. S8: Proliferation, homeostatic densities or crowding cannot explain winning of WT cells**. (**A**) Mitotic cells in mixed culture. Segmented nuclei (blue), outlines of E-cad KO clusters (green) and phospho-Histone H3 positive cells (pHH3, magenta) are shown. Right: Average fractions of mitotic cells in pure and mixed cultures. Each datapoint shows the fraction for one FOV. n=15 FOV from N=2 independent experiments (pure) and n=20, N=2 (mixed). (**B**) Cell densities in pure and mixed cultures over time. (**C**) Confocal images of WT and E-cad KO cells at homeostasis (4 days confluent culture). Right: Comparison of homeostatic densities to typical densities observed in cell competition (data already shown in Fig S2 A). Each datapoint shows the fraction for one FOV. n=11 (homeostasis) and n=16 (competition) FOV from N=2 independent experiments. P-value from unpaired t-test. (**D**) Left: Extrusion rates of MDCK cells over time. Rates are calculated for pure and mixed cultures. Right: Extrusion rates over cell densities, corresponding to (B). n=8 movies from N=2 (pure) and N=3 (mixed) independent experiments. P-values from Kruskal-Wallis test corrected for multiple comparisons (Dunn's test). Scale bars (A) 200 µm (B) 25 µm.

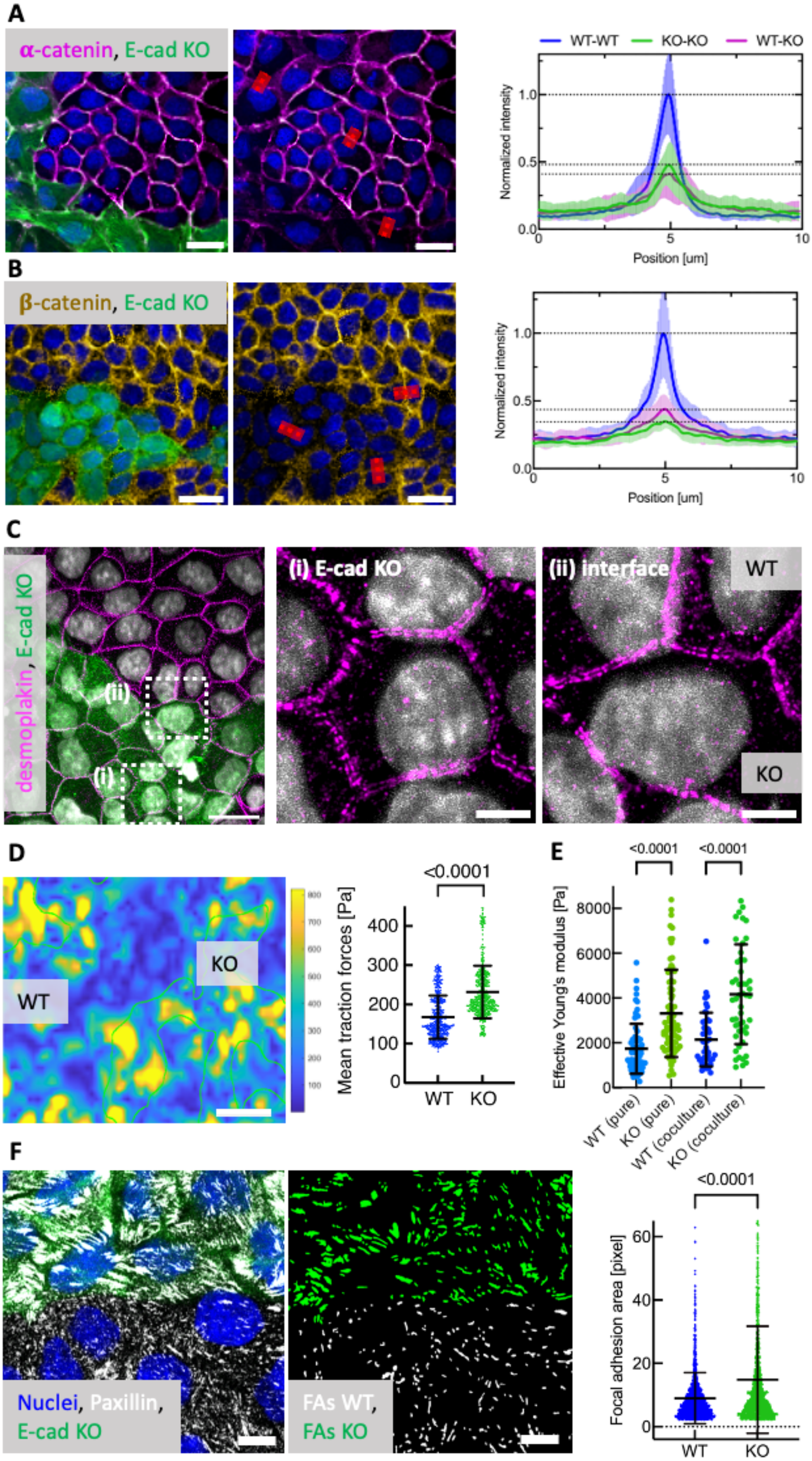
A
α-catenin, E-cad KO
WT-WT
KO-KO
WT-KO
Normalized intensity
Position [um]
B
β-catenin, E-cad KO
Normalized intensity
Position [um]
C
desmoplakin, E-cad KO
(i) E-cad KO
(ii) interface
WT
KO
D
WT
KO
Mean traction forces [Pa]
<0.0001
WT
KO
E
<0.0001
<0.0001
Effective Young's modulus [Pa]
WT (pure)
KO (pure)
WT (coculture)
KO (coculture)
F
Nuclei, Paxillin,
E-cad KO
FAs WT,
FAs KO
<0.0001
Focal adhesion area [pixel]
WT
KO

**Fig. S9: Differences in cell mechanics between MDCK WT and E-cad KO cells in mixed cultures.** E-cad KO cells express LifeAct-EGFP and are visualized in green in every image. (**A**) Confocal images showing accumulation of alpha-catenin (magenta) and (**B**) beta-catenin (yellow) at cell-cell junctions. Red lines indicate line plots. Right graphs are normalized to the highest average value. n=15 (alpha-catenin) and n=20 (beta-catenin) measurements from N=1 experiment. (**C**) Airyscan confocal images of desmoplakin showing desmosomes in the mixed cultures. Zoom-ins on (i) the bulk of E-cad KO cells and (ii) on the interface. (**D**) Color-coded traction force map on 15 kPa stiff surface. Green outline shows E-cad KO cluster. Right: Average traction forces. n=568 positions from N=4 independent experiments. (**E**) Cell stiffness measured by indentation for pure and mixed cultures. E-cad KO cells in mixed cultures were identified by their green fluorescence. n=46 (WT), n=51 (KO) indentations from N=2 independent experiments. (**F**) Confocal images showing Focal adhesions (Paxillin, white) and corresponding segmentation (middle). Right: Average FA area. n=1500 FAs representing N=2 independent experiments. P-values derived from unpaired t-test (D,F) or from Kruskal-Wallis test corrected for multiple comparisons (Dunn's test). All error bars show the standard deviation. Scale bars 50 µm (D); 20 µm (A, B, C); 10 µm (F); 5 µm (C, zoom-ins).

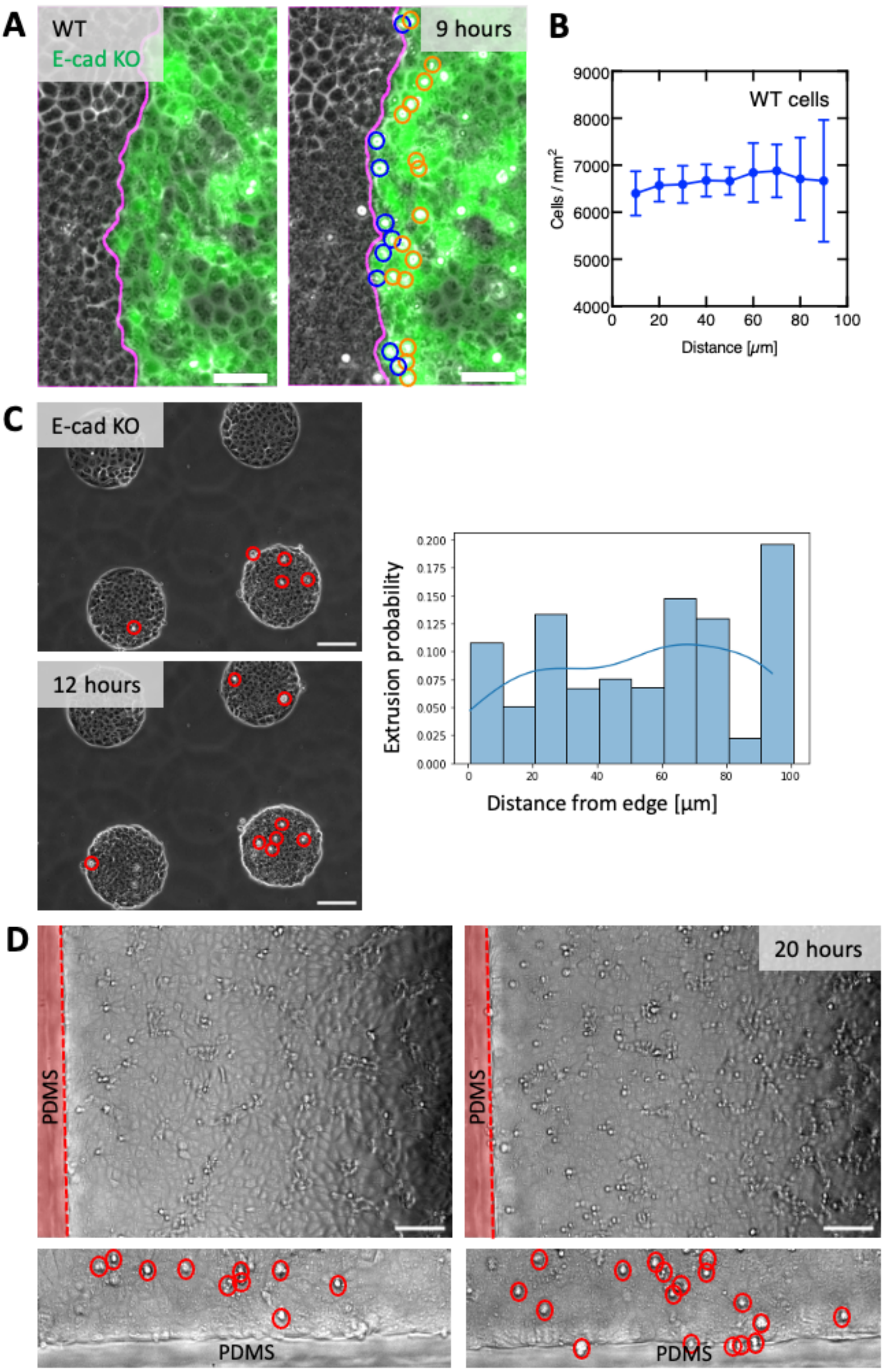


**Fig. S10: E-cad KO cells do not extrude at a passive interface.** (**A**) Example images of E-cad KO cells eliminated at the interface. Cells are eliminated directly in contact with WT cells (blue circles), or close to the interface without directly interacting with the WT cells (orange circles). Magenta line shows the interface. (**B**) Quantification of WT cell density over the distance to the interface based on nuclear segmentation shown earlier (Fig. S2 A). (**C**) Example images of E-cad KO cells confined on circular micropattern with a 100 µm radius. Red circles indicate extrusions. Below: Extrusion probability of confined E-cad KO cells. n=112 extrusions in N=1 experiment. (**D**) E-cad KO cells confined with a rigid passive fence (indicated by red shade, made of PDMS). Bottom: Zoom-ins on boundary area. Red circles indicate extrusions. No preferred accumulation of extrusions at the interface was observed. Scale bars 100 µm (D), 50 µm (A).

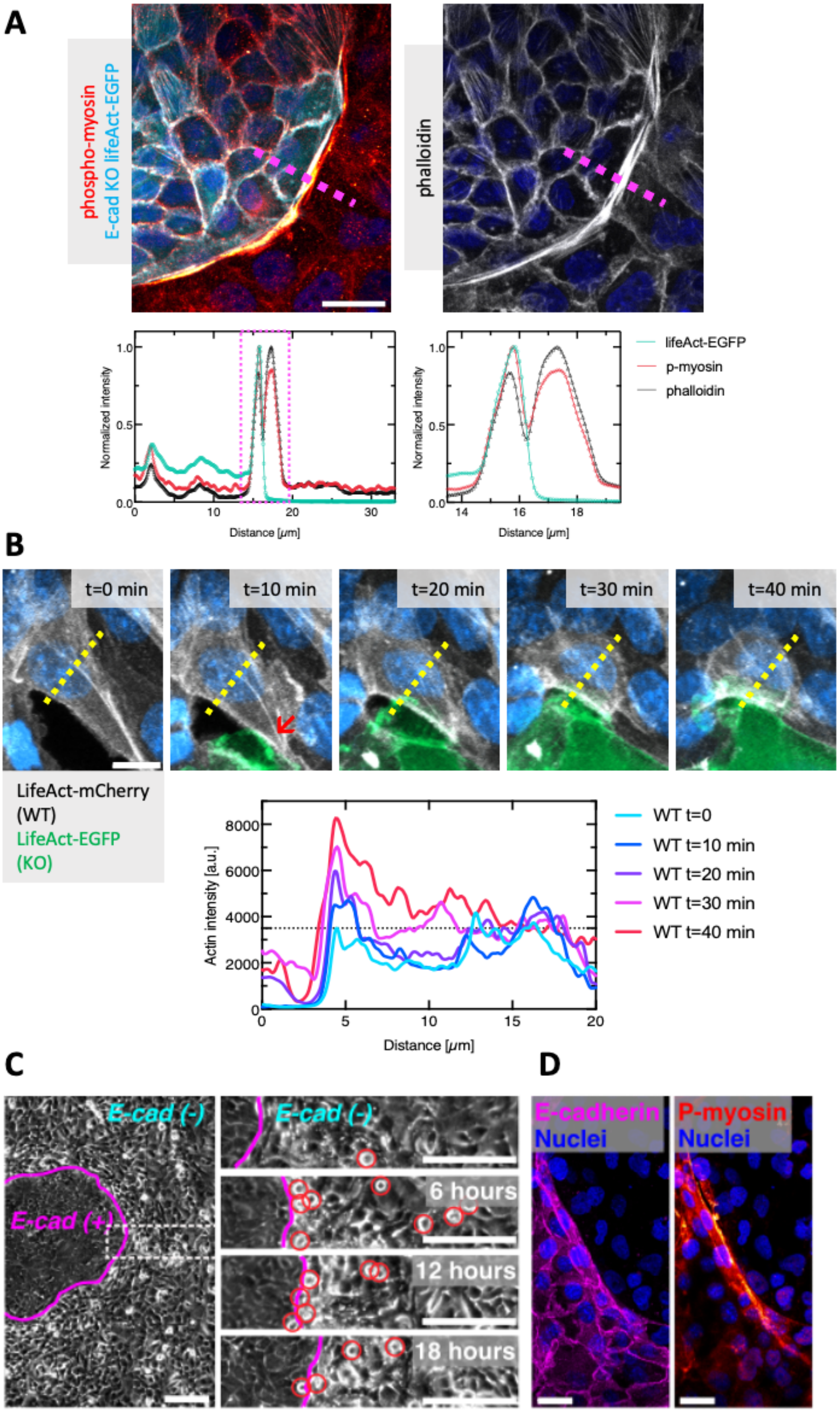
A
phospho-myosin
E-cad KO lifeAct-EGFP
phalloidin
Normalized intensity
Distance [μm]
lifeAct-EGFP
p-myosin
phalloidin
B
t=0 min
t=10 min
t=20 min
t=30 min
t=40 min
LifeAct-mCherry (WT)
LifeAct-EGFP (KO)
Actin intensity [a.u.]
Distance [μm]
WT t=0
WT t=10 min
WT t=20 min
WT t=30 min
WT t=40 min
C
E-cad (-)
E-cad (+)
E-cad (-)
6 hours
12 hours
18 hours
D
E-cadherin
Nuclei
P-myosin
Nuclei

**Fig. S11: Actomyosin accumulation at interface between E-cad KO and WT cells and interface elimination during intratumoral competition.** (**A**) Representative airyscan confocal image of increased phospho-myosin accumulating in both cell types at the interface. Intensity line plots were drawn at the indicated position for actin (phalloidin), phospho-myosin and LifeAct-EGFP (E-cad KO cells). Zoom-in on intensity profile reveals two separate intensity peaks in WT and E-cad KO and the colocalization of LifeAct-EGFP and phospho-myosin. (**B**) Actin dynamics following cell-cell collision. WT cells express LifeAct-mCherry (white) and E-cad KO cells express LifeAct-EGFP (green). Intensity values were measured at the indicated line. An increase in actin intensity within the WT cells was observed following collision (red arrow) at new interface with E-cad KO cells. (**C**) Representative phase-contrast image of patient-derived tumor xenograft cultured in 2D. E-cad (-) cells are eliminated at the interface. (**D**) Confocal images showing an increase of phospho-myosin at the tissue interface. Scale bars 200 µm (**C**), 20 µm (**A**), 25 µm (**D**), 10 µm (**B**).

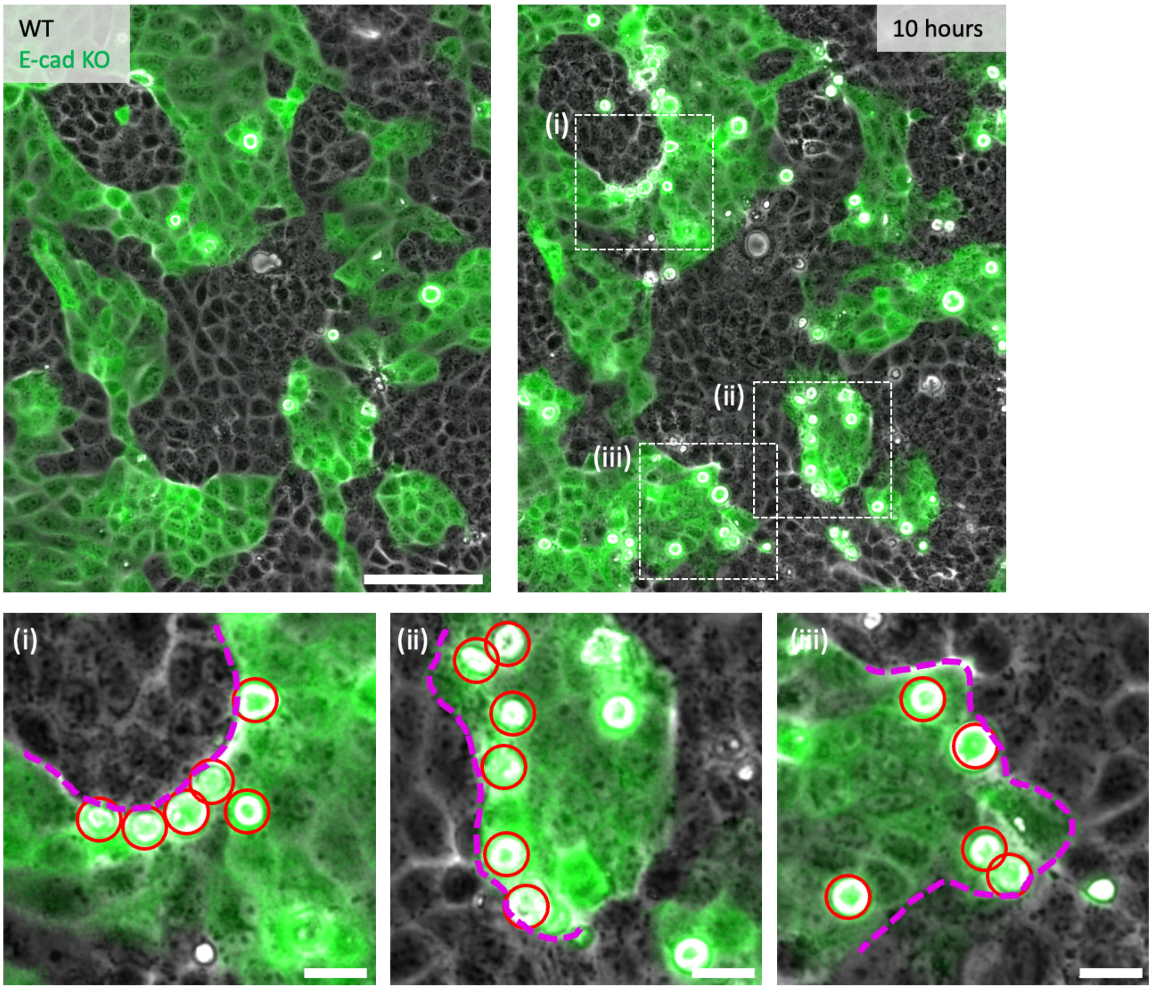


**Fig. S12: Preferred interface elimination is independent of interface curvature.** Brightfield and fluorescence (E-cad KO) images of a competing MDCK coculture. Within the same field of view, E-cad KO cells are eliminated at positively (i), minimally (ii) or negatively (iii) curved interfaces (dashed lines). Extrusions indicated by red circles. Scale bars: 100 µm (overview), 10 µm (zoom-ins).

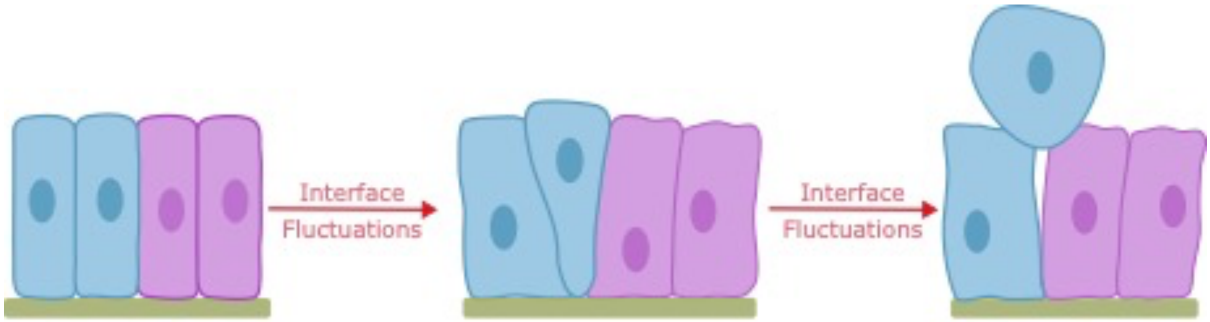


**Fig. S13: Schematic of shape changes.** The analytical model uses these three shapes: a cylinder, a cone and a sphere, to calculate the change in area used to find the work. All three cell shapes have the same volume.

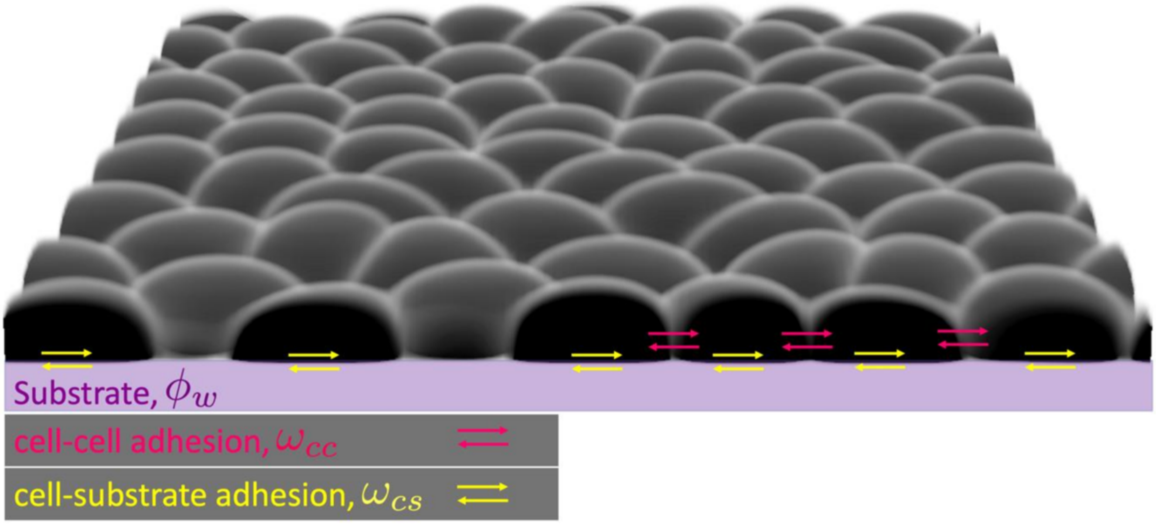


**Fig. S14: Key features of the multi-phase field model.** An example cross-section from a simulation. The arrows show, schematically, how cell-cell and cell-substrate adhesions are explicitly accounted for in the model and mathematically outlined in the Methods.

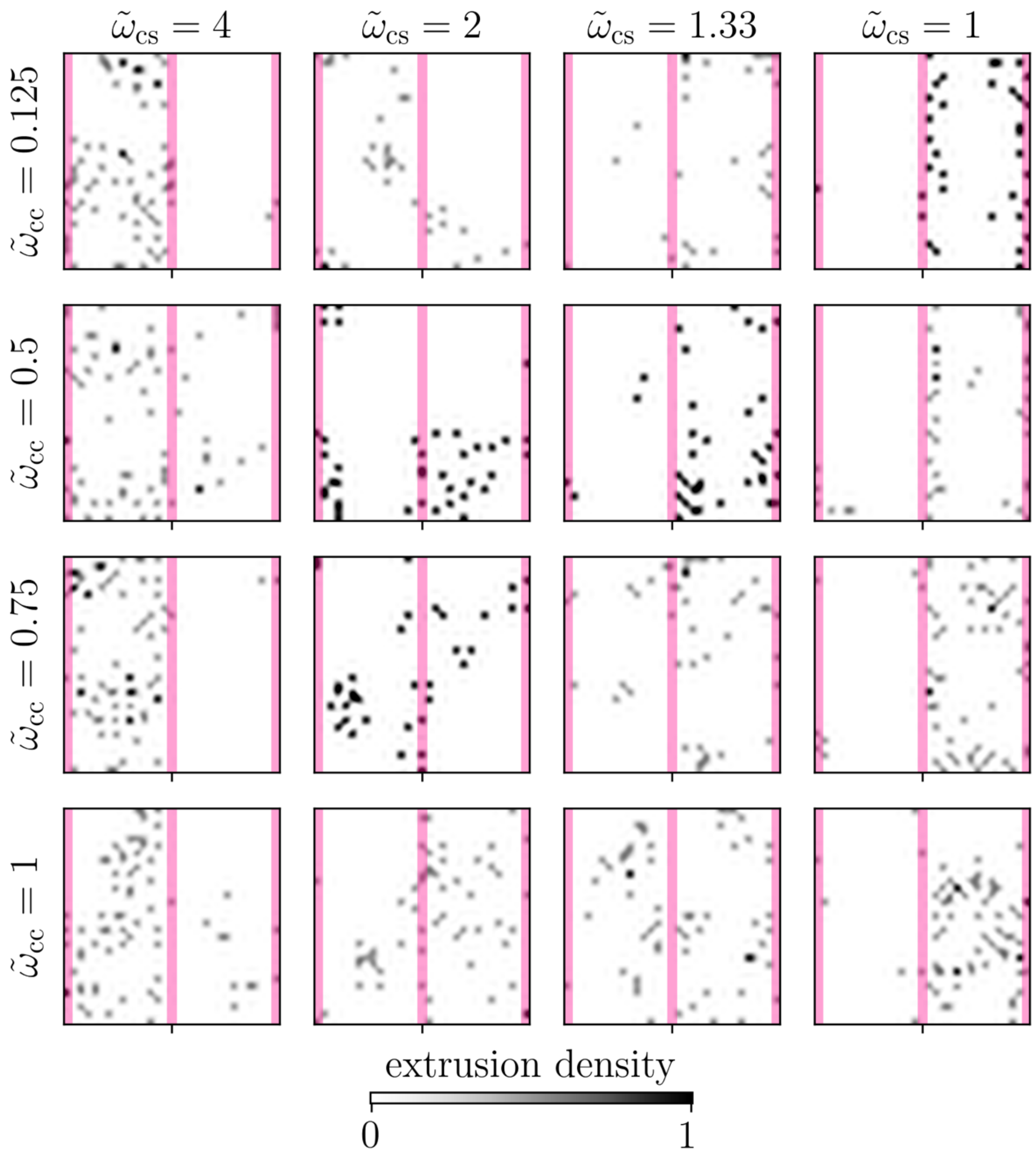


**Fig. S15: Regulation of type and location of winning cell sub-population by altering their mechanical interactions.** Two-dimensional extrusion density maps indicating the location of extrusions for various cell-substrate adhesion strengths $\widetilde{\omega}_{cs}$ and cell-cell adhesion strengths $\widetilde{\omega}_{cc}$. The red line indicates the domains associated with mWT cells (left side) and mE-cad KO cells (right side), keeping in mind the periodic boundary conditions.

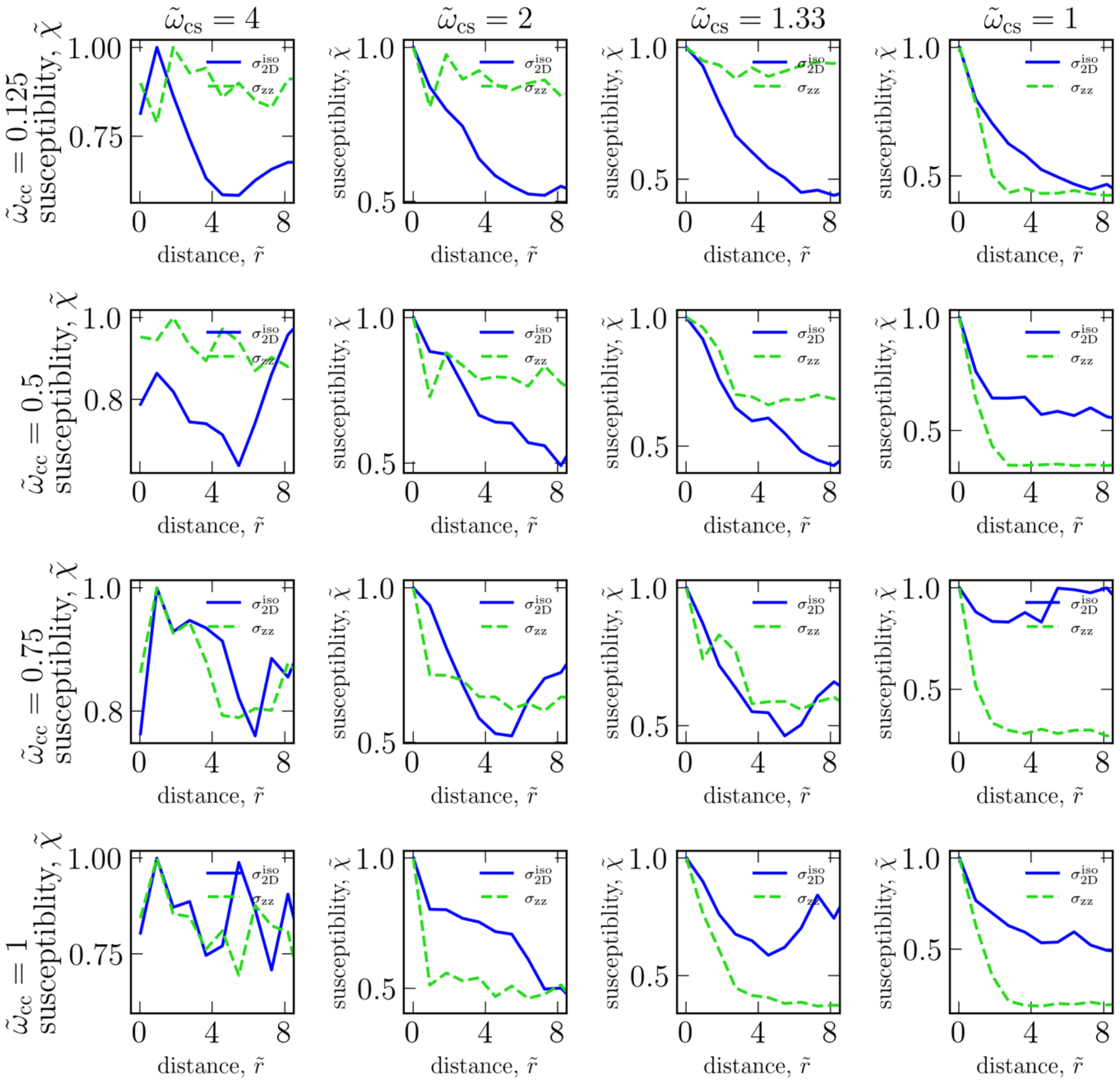


**Fig. S16: Stress fluctuations away from the interface highlight the role of stress transmission.** Susceptibility of two-dimensional isotropic stress ($\sigma_{2D}^{iso}$), and the out-of plane stress component ($\sigma_{zz}$) for mE-cad KO cells as a function of distance away from the interface for various cell-substrate adhesion ($\tilde{\omega}_{cs}$) and cell-cell adhesion ($\tilde{\omega}_{cc}$) strengths. The distance is normalized by the initialized cell radius. The susceptibility is normalized by the maximum value.

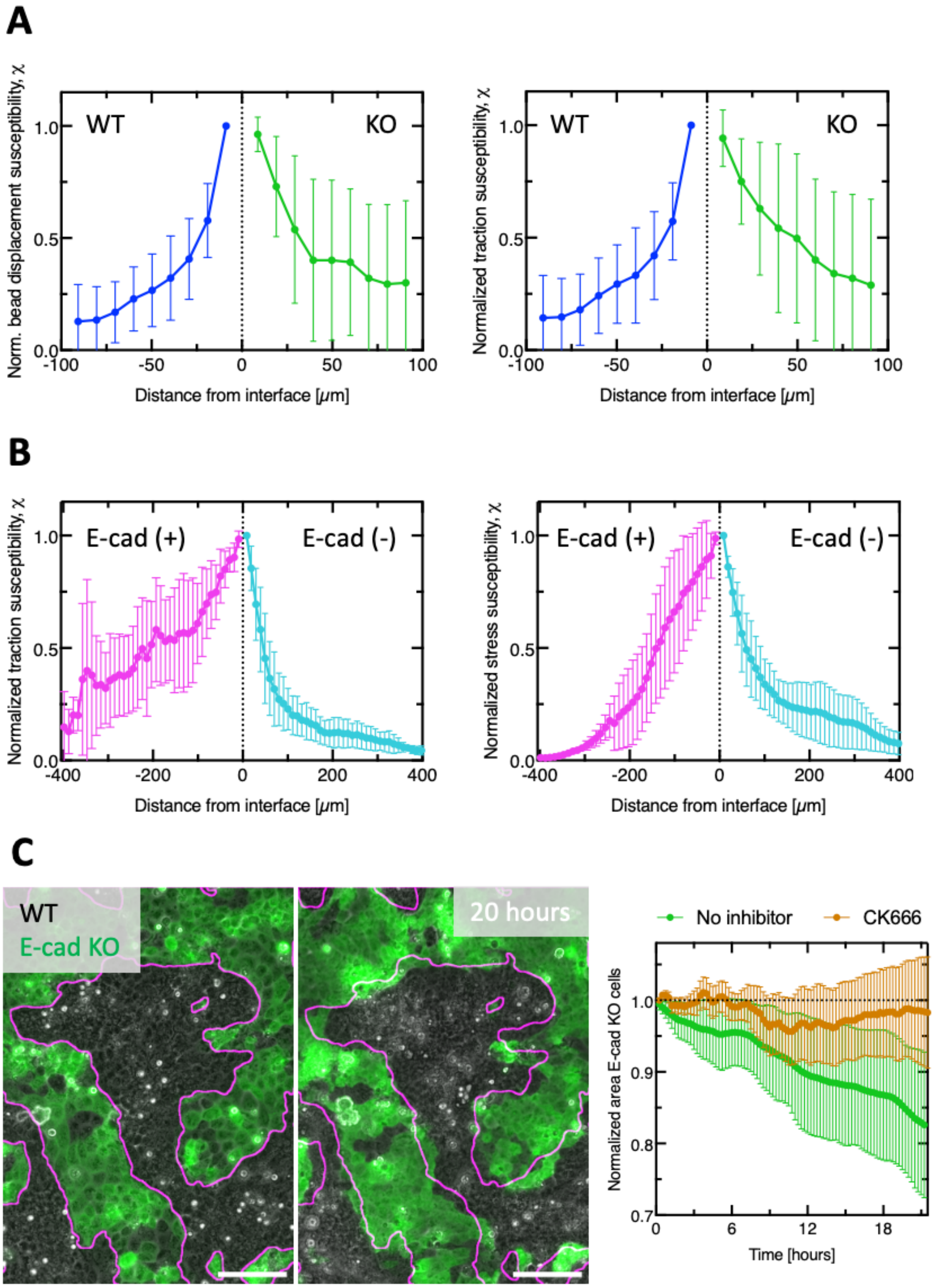


**Fig. S17: Interface fluctuations occur in MDCK cells and within patient-derived tumor xenograft.** (**A**) Susceptibility of bead displacement (left) and of traction forces (right) as a function of distance from the interface in competing MDCK WT and E-cad KO cells. (**B**) Susceptibility of tractions (left) and stresses (right) as a function of distance from the interface within patient-derived tumor xenograft cultured in 2D. Data from n=5 movies from N=2 independent experiments. (**C**) Area development of E-cad KO cells in competition with WT cells in the presence of 100 μM CK666, which inhibits protrusion formation. Means and standard deviations are shown. Values normalized to maximal value. Scale bars 100 μm.

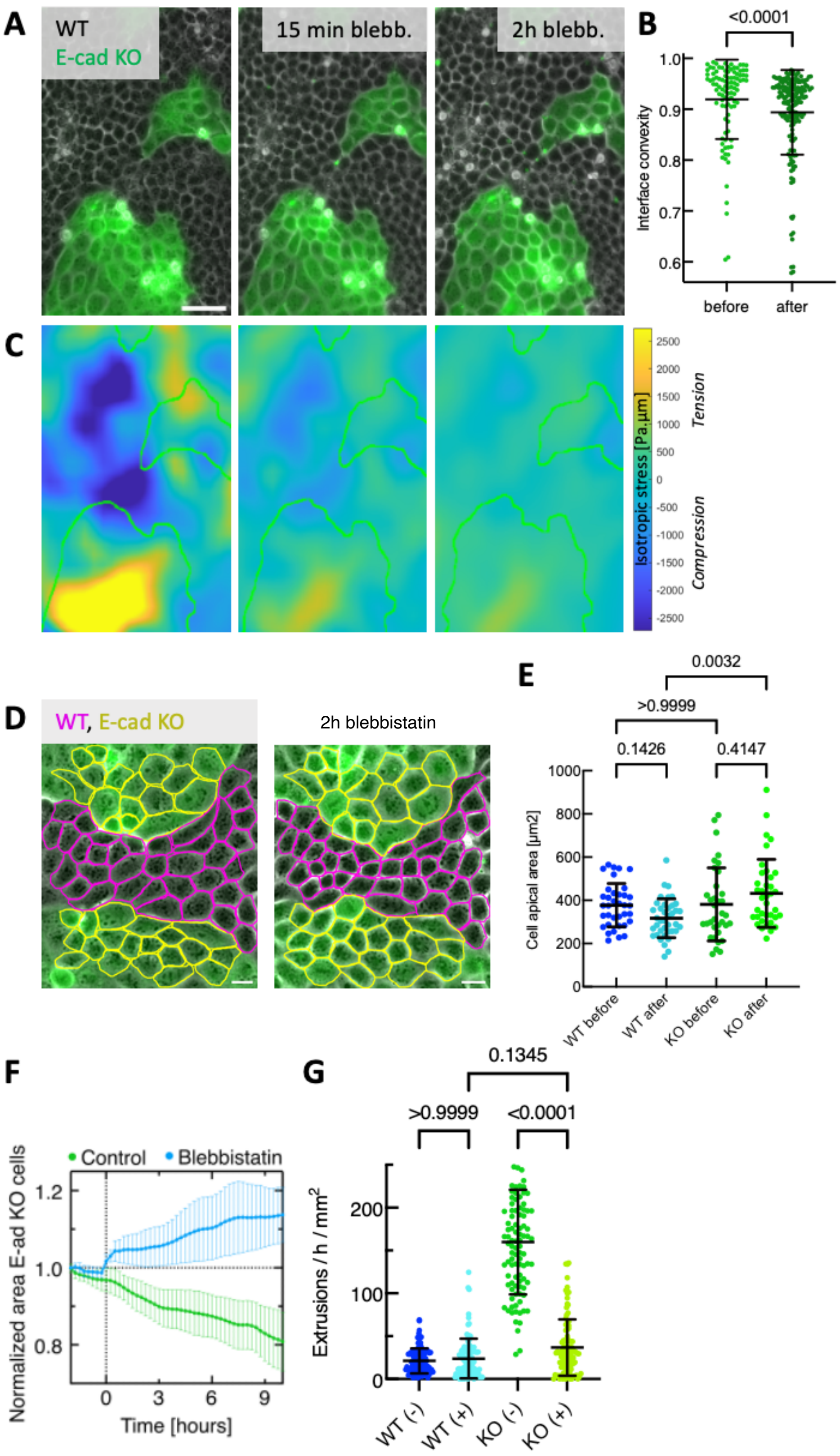
A
WT
E-cad KO
15 min blebb.
2h blebb.
B
<0.0001
Interface convexity
1.0
0.9
0.8
0.7
0.6
before
after
C
2500
2000
1500
1000
500
0
-500
-1000
-1500
-2000
-2500
Isotropic stress [Pa.µm]
Tension
Compression
D
WT, E-cad KO
2h blebbistatin
E
0.0032
>0.9999
0.1426
0.4147
Cell apical area [µm2]
1000
800
600
400
200
0
WT before
WT after
KO before
KO after
F
Control
Blebbistatin
Normalized area E-ad KO cells
1.2
1.0
0.8
0
3
6
9
Time [hours]
G
0.1345
>0.9999
<0.0001
Extrusions / h / mm$^2$
200
100
0
WT (-)
WT (+)
KO (-)
KO (+)

**Fig. S18: Effect of blebbistatin on cell competition between MDCK WT and E-cad KO cells.** (**A**) Additional representative images of the mixed cultures before and after the addition of 20 µM blebbistatin. (**B**) Quantification of the E-cad KO interface convexity before and after blebbistatin addition. Data from n=4 movies from N=2 independent experiments. (**C**) Isotropic stress fields before and after blebbistatin addition corresponding to images shown in (A). The red outline shows that WT remain under compression, the magenta outline shows that E-cad KO cells remain under tension. (**D**) Example pictures of apical areas before (left) and 2h after (right) blebbistatin addition. Apical areas were segmented manually. (**E**) Quantification of cell apical area change after 2h of blebbistatin addition. Every point shows a single cell. Data from N=1 independent experiment. (**F**) Area development of E-cad KO cells in competition with WT cells. Addition of blebbistatin at t=0 reduces interface fluctuations, which reverses the area development of E-cad KO cells. n=8 movies from N=2 independent experiments. (**G**) Comparison of extrusion rates after 20 µM blebbistatin addition compared to the standard condition. n=100 intervals from N=2 (+blebbistatin) and N=3 (-blebbistatin) independent experiments. P-values from unpaired t-test (B) or Kruskal-Wallis test corrected for multiple comparisons (Dunn's test) (E, G). Means and standard deviations are shown. Values normalized to maximal value. Scale bar 50 µm.

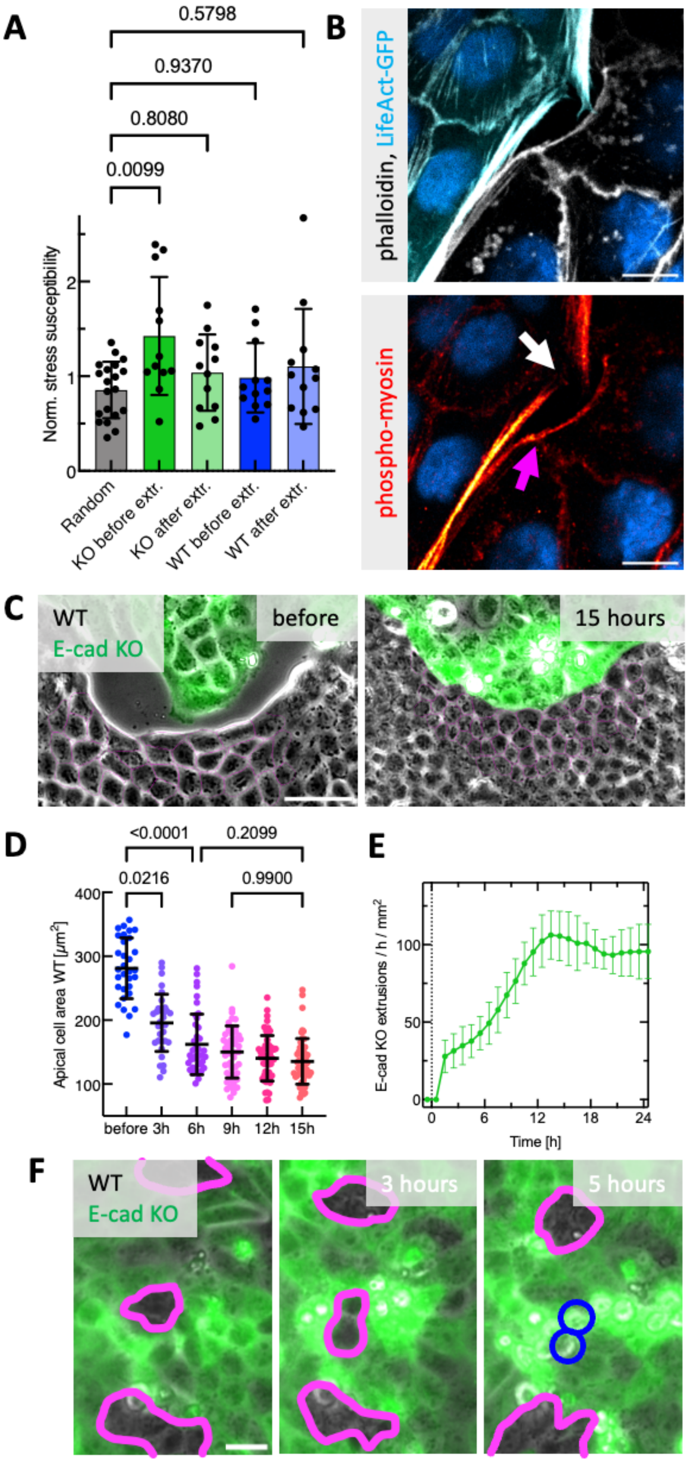


**Fig. S19:** (**A**) Comparison of the normalized stress susceptibility corresponding to Fig. 5h. Random positions of E-cad KO cells at the interface, E-cad KO and WT before (t=-40 and t=-20 in Fig. 5h) and after the extrusion event (t=0 and t=20) are shown. (**B**) Exemplary airyscan image of contractile actomyosin cables at the interface. Actin in white, phospho-myosin in red. Actomyosin fibers are interrupted between E-cad KO cells (white arrow) but can span multiple cells between WT cells (magenta arrow). (**C**) Phase contrast images of WT and E-cad KO cells colliding. Magenta outlines show manual area quantification of WT cells. (**D**) Manual segmentation of the WT apical area following tissue collision. Each datapoint represents one cell. Data from N=1 preparation. (**E**) Quantification of the extrusion rate over time. T=0 indicates the timepoint of collision. (**F**) Example of WT cell doublet being eliminated (blue circles) by surrounding E-cad KO cells. P-values from Kruskal-Wallis test corrected for multiple comparisons (Dunn's test). Means and standard deviations are shown. Values normalized to maximal value. Scale bar 50 µm (C), 20 µm (F), 10 µm (B).

**Video captions:**

**Video S1: Timelapse video of competition within patient-derived metaplastic cancer cells**. Phase contrast images of E-cad$^+$ cells surrounded by E-cad$^-$ cells corresponding to **Figure 1B**. Frame rate 10 min. Scale bar 200 µm.

**Video S2: Timelapse video of competition within patient-derived metaplastic cancer cells derived from a second patient**. Phase contrast images of E-cad$^+$ cells surrounded by E-cad$^-$ cells corresponding to **Fig. S1 C**. Frame rate 10 min. Scale bar 200 µm.

**Video S3: Timelapse video of competition between MDCK WT and MDCK E-cad KO cells**. Phase contrast images of E-cad KO cells expressing LifeAct-EGFP in green. Frame rate 15 min. Scale bar 100 µm.

**Video S4: Timelapse video of competition between MDCK dKO and MDCK E-cad KO cells**. Phase contrast images of E-cad KO cells expressing LifeAct-EGFP in green. Frame rate 15 min. Scale bar 100 µm.

**Video S5: Laser ablation of WT cluster within the mixed culture.** WT cells express CAAX-GFP and E-cad KO cells express LifeAct-EGFP. Video corresponds to Fig. S5 D. Magenta line shows ablation area. Frame rate 5 sec. Scale bar 50 µm.

**Video S6: Laser ablation of E-cad KO cluster within the mixed culture.** WT cells express CAAX-GFP and E-cad KO cells express LifeAct-EGFP. Video corresponds to Fig. S5 D. Magenta line shows ablation area. Frame rate 5 sec. Scale bar 50 µm.

**Video S7: Timelapse video of cell eliminations at the interface between MDCK WT and MDCK E-cad KO cells.** Phase contrast images of E-cad KO expressing LifeAct-EGFP in green. Magenta line highlights the interface. Note that E-cad KO cells are not only eliminated when directly in contact with WT cells. Frame rate 15 min. Scale bar 50 µm.

**Video S8: Example video showing simulation of competition between model WT and model E-cad KO cells.** mWT cells are shown in blue, mE-cad KO cells in red. The cell-cell adhesion strength of mWT cells is 8x higher than the mE-cad KO, the cell-substrate adhesion strength is constant.

**Video S9: Timelapse video of actin dynamics in competition between MDCK WT and E-cad KO cells**. Confocal images showing maximum projection of actin expressed in both cell types. 60% MDCK WT cells express LifeAct-mRuby, displayed in white, all E-cad KO cells express LifeAct-EGFP, displayed in green. Note the protrusion activity of E-cad KO cells at the interface. Frame rate 7.5 min. Scale bar 20 µm.
